# VGLL3-centered network connects placental, vascular, and immune defects in preeclampsia

**DOI:** 10.1101/2025.05.30.657097

**Authors:** Olesya Plazyo, Laura B Chopp, Rishyanth Peela, Kelly Young, Haihan Zhang, Rachael Bogle, Ashley Hesson, Elizabeth S Langen, Ingrid L Bergin, Li-Jyun Syu, Jake Erba, Joseph Kirma, Poulami Dey, Lin Zhang, Mrinal K Sarkar, William R Swindell, Katherine A Gallagher, Nicole L Ward, Kanakadurga Singer, J Michelle Kahlenberg, Allison C Billi, Andrzej A Dlugosz, Santhi K Ganesh, Lam C Tsoi, Johann E Gudjonsson

## Abstract

Preeclampsia affects approximately 1 in 10 pregnancies, leading to severe complications and long-term health risks for both mother and offspring. While the etiology remains unclear, preeclampsia has been linked to both autoimmunity and the timing of menarche. Through human single-cell and spatial analyses, coupled with in vitro, in vivo, and ex vivo models, we demonstrate that VGLL3, a transcription co-regulator in the Hippo pathway, is upregulated in preeclamptic placentas. VGLL3 promotes immune activation, impairs trophoblast differentiation, and induces endothelial dysfunction, all of which contribute to pregnancy-related hypertension, fetal growth restriction, and offspring mortality. Our data reveal that VGLL3 acts upstream of preeclampsia-associated processes, including the production of sFLT1, a key biomarker of the disease. Notably, targeting VGLL3—either by genetic deletion in mouse placentas or through therapeutic inhibition in human placentas—protects against preeclampsia and alleviates disease pathology. These findings position VGLL3 as a promising novel therapeutic target for preeclampsia.

## INTRODUCTION

Preeclampsia (PreE) is a vascular endothelial disorder of pregnancy accompanied by hypertension and associated with severe perinatal morbidity for mother and child, including death and lifelong complications (1). Due to the limited understanding of PreE pathogenesis, FDA-approved treatment options remain scarce, with delivery of the fetus and placenta serving as the only definitive intervention.

Previous studies have broadly categorized the major pathological mechanisms underlying PreE into four key areas: placental dysfunction (including inadequate spiral artery remodeling and a shift toward anti-angiogenic signaling), metabolic disturbances, maternal anti-fetal immune rejection, and dysregulation of extracellular matrix (ECM) proteins (2). A hallmark of PreE is increased placental secretion of soluble vascular endothelial growth factor receptor 1 (sFLT1/VEGFR1) (3) and reduced production of placental growth factor (PlGF) (4). The ratio of these two protein markers currently forms the basis of the only FDA-approved test for assessing PreE risk (4). Although therapeutic strategies targeting sFLT1 have shown promise in preclinical animal models (5), their effectiveness in human patients remains under investigation. Deciphering the complex etiology of PreE is essential for identifying additional therapeutic targets.

In here, by integrating single-cell and spatial transcriptomic analyses with human placental explants, mouse models, and *in vitro* systems, we demonstrate that hallmarks molecular features of PreE center around a VGLL3-centered network linking together the major pathological mechanisms in PreE and the well-known association between PreE and autoimmunity (6, 7) and early-age of menarche (8, 9).

## RESULTS

### Single-cell and spatial RNA seq revealed activation of immune cells, altered signaling in non-immune cells, and Hippo pathway dysregulation in PreE placenta

To characterize the cellular composition and transcriptomic alterations in placentas affected by preeclampsia (PreE), we performed single-cell RNA sequencing (scRNA-seq) on placental tissue from three healthy donors (HD) and eight individuals with PreE (Table 1). After rigorous quality control and filtering, the dataset comprised 45,813 high-quality cells, with an average of 10,664 transcripts and 2,560 genes per cell. Data preprocessing using Seurat and integration with Harmony (10) enabled unsupervised clustering based on differentially expressed genes (DEGs), resulting in the identification of 23 distinct cell clusters (Fig. 1a). Visualization using Uniform Manifold Approximation and Projection (UMAP) and cluster annotation based on cluster- and lineage-defining genes (Sup. Table 1; Sup. Fig. 1, 2a–b) revealed six major placental cell types: trophoblasts, maternal-derived stromal cells, endothelial cells (ECs), lymphocytes, and myeloid lineage cells (Fig. 1a, left). While the overall composition of immune and non-immune clusters was comparable between HD and PreE placentas, we observed an increased proportion of monocytes, B cells, and extravillous trophoblasts (EVTs), alongside a decreased proportion of a macrophage subpopulation (Mac3) and αβ T cells in PreE samples (Fig. 1a, right). Pseudobulk analysis across all cell populations confirmed significant upregulation of PreE-associated genes in the PreE cohort (Fig. 1b, left). Ingenuity Pathway Analysis (IPA) of predicted upstream regulators driving the PreE transcriptomic signature highlighted several key inflammatory mediators, including TNF, TGFβ1, IFNγ, IL-1B, IL-4, and IL-33 (Fig. 1b, right), reinforcing the central role of immune dysregulation in PreE pathogenesis.

**Fig. 1.**
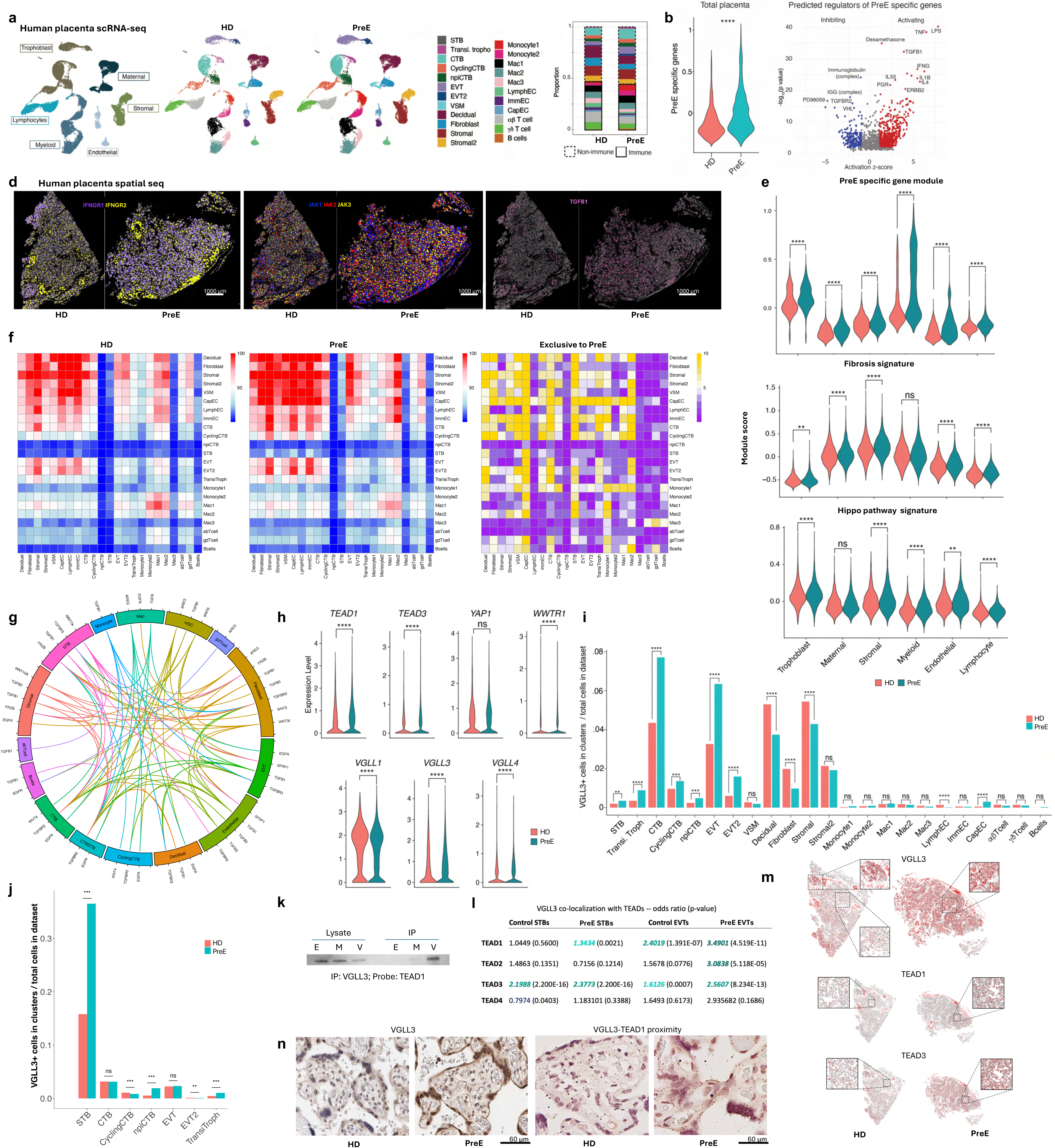
scRNA and spatial RNA seq implicate Hippo pathway in PreE. a. UMAP plot showing placental cells from healthy donors (HD) or patients with PreE colored by main cell types (left) and their sub-types (center). Barplot showing proportion of clusters in each condition (right, same color coding). b. Violin plot shows module score of PreE specific genes calculated on total placenta between conditions. Significance calculated with a Wilcoxon Rank Sum test. **** p < 0.0001 (left). c. Volcano plot shows putative regulators of this gene signature predicted by Ingenuity Pathway Analysis (Activation z-score [x-axis] vs. P-value, −Log10 scale [y-axis]). Red dots indicate activating regulator, blue dots indicate repressive regulator. Volcano plots show differential gene expression (Log2 fold-change [x-axis] vs. P-value, −Log10 scale [y-axis]) between HD and PreE in indicated myeloid clusters. d. Expression patterns of IFNG receptors (left), JAK kinases (center), and TGFB1 (right) determined by spatial seq. e. Gene module scores were calculated to assess PreE-, fibrosis-, and Hippo pathway-specific gene expression across main cell populations. f. Ligand-receptor interaction analyses in healthy and PreE placentas allowed identification of PreE-specific cell-cell interactions. g. Circos plot of PreE-specific Hippo pathway receptor-ligand interactions in major cell types. h. Violin plots showing expression of main regulators in Hippo pathway in HD and PreE placentas. i. Bar plot shows proportion of VGLL3+ expressing cells in individual clusters over total cells in the sample based on scRNA-Seq data. Significance calculated with 2-sample test for equality of proportions with continuity correction. j. Bar plot shows proportion of VGLL3+ expressing cells in individual clusters over total cells in the sample based on spatial seq data. k. Co-IP confirmed VGLL3-TEAD1 binding. HTR8 cells were transfected either with empty vector (E), mutated VGLL3 (M), or full-length WT VGLL3 (V). l. Co-localization of VGLL3 with TEADs in STBs and EVTs computed from Xenium spatial seq. Odds ratios and p-values are shown. m. Expression pattern of VGLL3, TEAD1, and TEAD3 determined by spatial seq. n. IHC (left) confirmed increased expression of VGLL3 on the protein level in PreE placentas. Increased VGLL3-TEAD1 proximity in PreE placentas was confirmed by PLA (right).

**Table 1.** Demographics of donors whose placentas were used for scRNA-seq analyses.

|  | **HD (n=3)** | **PreE (n=8)** | **P-value** |
| --- | --- | --- | --- |
| Maternal age (years)<sup>a</sup> | 31.7 (21-38) | 32.6 (23-41) | >0.9999 |
| Race<sup>b</sup> | White=66.7% (2/3)<br>Black=33.3% (1/3) | White=75% (6/8)<br>Black=12.5% (1/8)<br>Hispanic=12.5% (1/8) | >0.9999<br>0.4909<br>>0.9999 |
| Nulliparity<sup>b</sup> | 33.3% (1/3) | 50% (4/8) | >0.9999 |
| Gestational age at delivery (days)<sup>a</sup> | 277.3 (273-285) | 267 (251-284) | 0.2605 |
| Gestational age at diagnosis (days)<sup>a</sup> | N/A | 262.9 (249-284) | N/A |
| Chronic hypertension (cHTN) | 0% (0/3) | 0% (0/8) | N/A |
| Incidence of PreE with severe features | N/A | 50% (4/8) | N/A |
| Incidence of autoimmune disease<sup>b</sup> | 0% (0/3) | 12.5% (1/8) | >0.9999 |
| Incidence of labor<sup>b</sup> | 66.7% (2/3) | 75% (6/8) | >0.9999 |
| Cesarean delivery<sup>b</sup> | 33.3% (1/3) | 62.5% (5/8) | 0.5455 |

Data are shown as mean (range) and percentage (n/N). Statistics analyzed by <sup>a</sup>Mann-Whitney U-test or <sup>b</sup>Fisher's exact test.

Extensive evidence supports a role for immune dysregulation in the pathogenesis of PreE (11), a finding further substantiated by our single-cell transcriptomic data. Monocytes and macrophages from PreE placentas exhibited elevated expression of inflammatory genes, including interferon-responsive genes as well as *CXCL2*, *IL1A*, *CCL3*, and *CCL20* (Sup. Fig. 2a)—all associated with immune activation, endothelial dysfunction, and previously implicated in PreE pathophysiology (12–15). In addition, T cell subsets in PreE placentas showed increased expression of inflammatory mediators: *IL32* and *GZMA* were enriched in γδ T cells, while CD8 T cells upregulated *IFNG*, *GZMH*, and *GZMA* (Sup. Fig. 2b). CD4 T cells in PreE placentas displayed enhanced expression of signaling molecules such as *FYN* and *JUNB*, but reduced expression of functional molecules *CD40LG* and *TNF*, consistent with reported impairments in regulatory T cell function in PreE (16).

Although B cells represented a relatively small fraction of the total placental immune population, they demonstrated notable gene expression changes in PreE. Specifically, B cells from PreE placentas showed reduced expression of *SPIB*, a transcription factor that limits plasma cell differentiation, along with increased expression of genes involved in immunoglobulin production and plasma cell function, including *PRDM1*, *CD27*, *MZB1*, and *XBP1* (Sup. Fig. 2c). Pathway analysis of these B cells revealed enrichment of IL-12-mediated signaling cascades that drive a B cell–intrinsic, IFNγ-dependent feed-forward loop promoting plasmablast differentiation (Sup. Fig. 2d) (17). Additionally, expression of *CXCR4*, a receptor for stromal-derived factor 1 (SDF-1) that supports B cell survival under hypoxic conditions, was elevated, suggesting a potential adaptation to the altered placental microenvironment in PreE (18).

To assess spatial transcriptomic differences between HD and PreE placentas, we utilized Xenium spatial profiling on placental sections encompassing both maternal (decidua) and fetal (chorionic villi) compartments, identified by H&E and HLA-G positivity (Sup. Fig. 3a). By overlaying cell-type markers identified through our scRNA-seq dataset, we successfully mapped all 23 identified cell types across the tissue architecture. Notably, single-nucleus sequencing enabled improved detection of multinucleated syncytiotrophoblasts (STBs) (Sup. Fig. 3b).

Spatial transcriptomic analysis confirmed the presence of immune dysregulation in PreE and revealed compartment-specific expression patterns of key signaling molecules. For example, while both *IFNGR1* and *IFNGR2*, the receptors for interferon-γ (IFNγ), were upregulated in PreE placentas, their spatial distribution diverged: *IFNGR1* was predominantly localized to the fetal villous compartment, whereas *IFNGR2* was confined to the maternal decidua (Fig. 1d). These findings underscore the importance of spatial context in understanding the compartmentalized immune responses associated with PreE.

Strikingly, trophoblasts, stromal cells, and other non-immune cell types exhibited a robust increase in PreE gene module scores, comparable to those observed in immune cells (Fig. 1e, top). Receptor-ligand interaction analysis predicted numerous PreE-specific signaling events between non-immune cell populations (Fig. 1f). Further analysis of complement and cytokine receptor-ligand expression patterns (Sup. Fig. 4a–b) revealed distinct contributors to the inflammatory microenvironment in PreE, notably identifying decidual cells as a key source of *C3* and *CCL28*. Gene expression changes in PreE non-immune cells (Sup. Fig. 5a) implicated multiple dysregulated pathways, including networks centered on HIF-1α and AP-1 transcription factors, Syndecan-1– and focal adhesion kinase (FAK)–mediated signaling, and β1 integrin cell-surface interactions (Sup. Fig. 5b). In addition, fibrosis module scores were significantly elevated across most non-immune cell types, suggesting broad engagement of fibrotic processes in the PreE placenta (Fig. 1e, middle).

Further analysis identified dysregulation of the Hippo signaling pathway—a known driver of fibrosis in various pathological contexts (19)—as a prominent feature of PreE (Fig. 1e, bottom). Receptor-ligand interaction analysis of Hippo pathway–associated genes revealed robust intercellular communication involving both immune and non-immune cell populations (Fig. 1g). We next assessed the expression of key Hippo pathway effector genes and observed significant upregulation of multiple *TEAD* family transcription factors and *VGLL* family transcription co-regulators, but not *YAP1*, in PreE placentas (Fig. 1h, Sup. Fig. 6a-b). VGLL proteins lack DNA-binding domains and regulate gene expression by binding TEAD transcription factors. Given the established role of *VGLL3* in autoimmune disease (20, 21), we focused on its expression across placental cell types and observed a markedly increased proportion of *VGLL3*-expressing cells among PreE trophoblasts (Fig. 1i–j).

Co-immunoprecipitation and subsequent mass spectrometry confirmed physical interactions between VGLL3 and TEAD1/TEAD3 (Fig. 1k and Sup. Table 2). Spatial transcriptomic analysis further validated co-expression of VGLL3 with TEAD1 and TEAD3 in STBs and EVTs, with significantly higher co-localization scores and odds ratios in PreE (Fig. 1l). In healthy placentas, VGLL3^hi trophoblasts were restricted to discrete regions, predominantly near the decidua, whereas in PreE placentas, they expanded across broader placental regions (Fig. 1m)—a distribution pattern paralleled by *FLT1* expression (Sup. Fig. 6c). Finally, proximity ligation assay (PLA) confirmed increased physical proximity between VGLL3 and TEAD1 in PreE placentas, supporting their interaction *in vivo* (Fig. 1n).

To validate our findings in a larger and independent cohort, we reanalyzed our previously published bulk RNA-seq data derived from a substantial number (n=84) of formalin-fixed paraffin-embedded placental samples (22). Despite the lower resolution of bulk sequencing compared to single-cell approaches, the results showed strong concordance with our scRNA-seq data. Differentially expressed genes (DEGs) in PreE placentas included *IGFBP1*, a gene with polymorphisms previously associated with increased PreE risk (23), and *DDX20*, a regulator of miRNA biogenesis in trophoblasts (24) critical for embryonic development (25) (Sup. Fig. 7a). Among the top dysregulated pathways were MHC class II, Smad, and TGF-β signaling (Sup. Fig. 7b), all of which are known to play roles in immune modulation and placental development.

IPA further identified *VGLL3* as a predicted upstream regulator of the bulk RNA seq PreE gene expression signature (Sup. Fig. 7c). As a transcriptional co-regulator, VGLL3 is poised to influence multiple signaling cascades simultaneously by modulating the expression of diverse target genes. Thus, VGLL3 dysregulation in PreE likely contributes to multiple pathogenic processes in parallel. To further dissect the cellular and molecular consequences of VGLL3 activation in PreE, we next integrated our single-cell and spatial transcriptomics datasets to characterize VGLL3-driven shifts in placental biology.

### VGLL3 upregulation in PreE trophoblasts promotes inflammatory responses and apoptosis in STBs and abnormal interaction of EVTs with immunomodulatory ECs

The Hippo signaling pathway has been implicated in regulating the stemness and differentiation of placental trophoblasts (26). To assess how TEAD1 and VGLL3 expression changes during trophoblast differentiation, we applied Monocle3 to our scRNA-seq data to infer developmental trajectories in trophoblast cells from both HD and PreE placentas (27). While the overall pseudotime trajectories were comparable between conditions, TEAD1 and VGLL3 expression steadily increased across pseudotime in PreE extravillous trophoblast (EVT) cells, in contrast to a rise-and-fall pattern observed in HD samples (Fig. 2a).

**Fig. 2.**
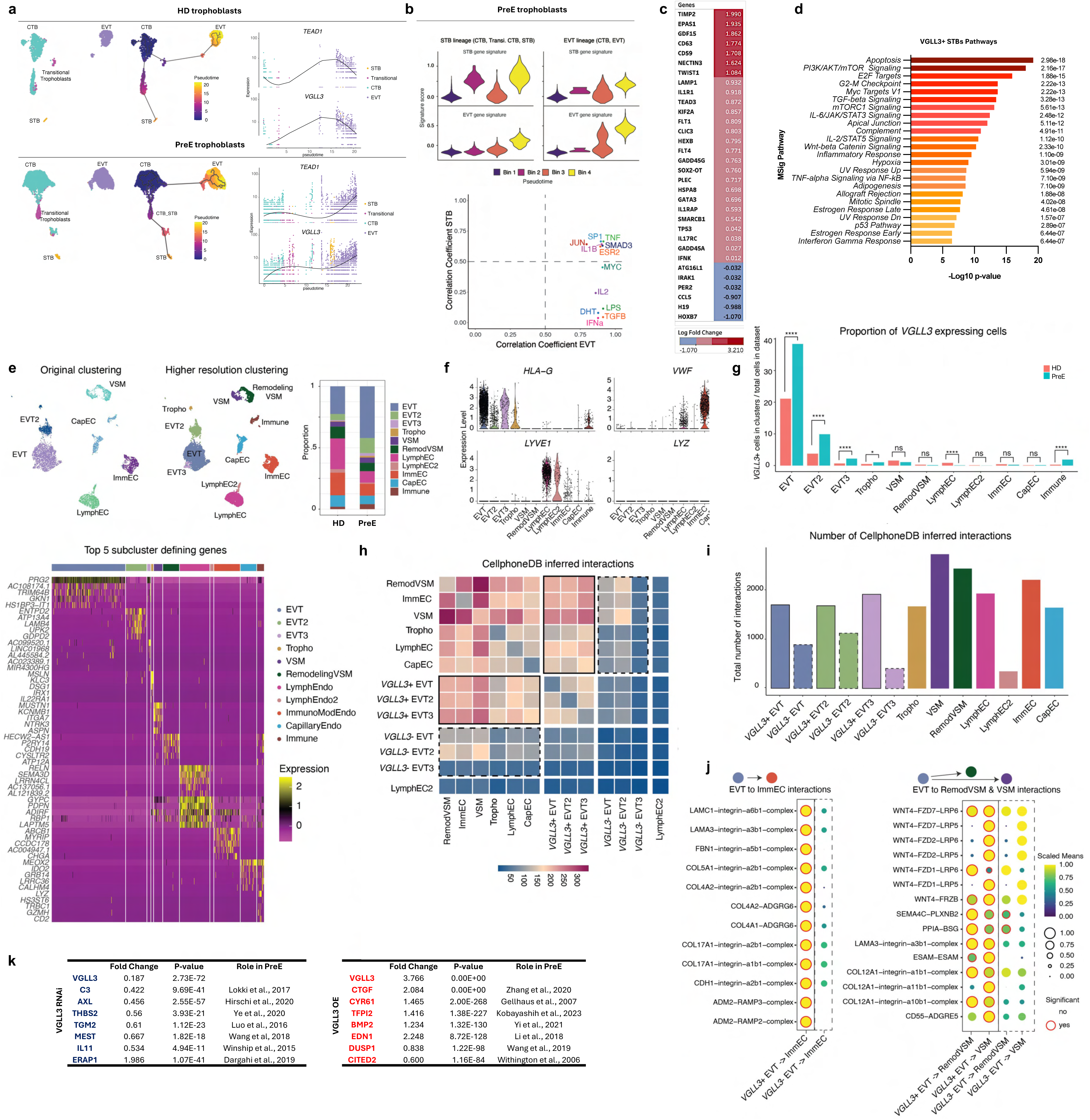
Aberrant VGLL3 expression in PreE trophoblasts promotes inflammatory responses and apoptosis in STBs and abnormal interaction of EVTs with immunomodulatory ECs. a. Trophoblast cells are shown on UMAP plots generated by Monocle3-derived dimensional reduction, and colored-coded by Seurat-defined clusters (left) or pseudotime value (right). Thick lines (right) are developmental trajectories defined by pseudotime analysis. Gene expression plots show expression of indicated genes across pseudotime, color coded by Seurat-defined clusters. b. (Top) Violin plots show module score of gene expression signatures across pseudotime bins. Signatures were defined by identifying genes that increased in trophoblast lineages across pseudotime that were also higher in PreE compared to HD. IPA was used to identify putative upstream regulators of these signatures. (Bottom) Scatter plot showing the correlation coefficients between the module target score of putative upstream regulators and EVT pseudotime (x*-*axis) and STB pseudotime (y*-*axis). c. Select genes differentially expressed in VGLL3+ STB (pval < 0.05). d. Pathways dysregulated in VGLL3+ STBs vs VGLL3-STBs based on MSigDB Hallmark 2020 database. e. UMAP plots showing EVT, endothelial, and VSM cells from human dataset colored by original cluster (top left) or sub-cluster (top center). Bar graph shows proportions of cells within conditions (top right). Heatmap lists cluster defining genes (bottom). f. Violin plots show expression of indicated genes across subclusters. g. Bar plot shows proportion of VGLL3+ expressing cells in individual subclusters over total cells in the sample. Significance calculated with 2-sample test for equality of proportions with continuity correction. * p < 0.05, ** p< 0.01, *** p < 0.001,**** p < 0.0001. h. CellphoneDB was used to identify inferred ligand-receptor interactions between VGLL3+ and VGLL3-trophoblast populations and endothelial or VSM cells from pre-eclamptic placental tissue. Heatmap showing number of ligand-receptor pairs between indicated populations. Solid line indicates VGLL3+ cells, dashed line indicates VGLL3-cells. i. Number of inferred interactions between indicated populations. Solid line indicates VGLL3+ cells, dashed line indicates VGLL3-cells. j. Dot plots show scaled mean scores of indicated interactions between VGLL3+ and VGLL3-EVT and ImmEC cells (left) or EVT and VSM cells (right). k. Select PreE-associated DEGs after VGLL3 knock-down (RNAi, top) in HTR8 cells or overexpression (OE, bottom) in HEK293 cells.

To identify potential regulators of the PreE-specific gene expression changes during trophoblast differentiation, we first defined gene signatures comprising transcripts that were both upregulated across pseudotime in PreE EVTs or STBs and preferentially expressed in PreE over HD cells (Fig. 2b, top). We then used IPA to infer upstream regulators of these signatures. For each predicted regulator, we calculated a module score based on its downstream target genes and assessed its correlation with pseudotime. This analysis revealed several inflammatory drivers—including lipopolysaccharide (LPS), type I interferons, IL-2, and TNF—as candidate regulators of the PreE-associated transcriptional shift (Fig. 2b, bottom). Together, these findings suggest that the trajectory of VGLL3 and TEAD1 expression is dysregulated during trophoblast differentiation in PreE and is accompanied by a progressive, pro-inflammatory transcriptional program in both STBs and EVTs.

To further characterize the functional differences between VGLL3⁺ and VGLL3⁻ STBs, we interrogated our spatial transcriptomics data. VGLL3⁺ STBs showed differential expression of several genes previously implicated in PreE, including TIMP2, TWIST1, and FLT1 (Fig. 2c). Pathway enrichment analysis revealed that VGLL3⁺ STBs were associated with activation of key signaling pathways such as apoptosis, PI3K/AKT/mTOR, TGF-β, and IL-6/JAK/STAT3 (Fig. 2d). These findings are particularly relevant given that STBs in PreE are known to exhibit increased apoptosis, altered nutrient transport, and enhanced shedding of proinflammatory mediators— processes that contribute to the formation of syncytial knots and systemic maternal inflammation. The transcriptional profile of VGLL3⁺ STBs suggests that VGLL3 may act upstream of these dysregulated processes in PreE, serving as a potential driver of STB dysfunction.

Successful pregnancy depends on the ability of EVTs to invade the maternal decidua and remodel spiral arteries, a process that involves the replacement of vascular smooth muscle (VSM) cells and ECs. Impaired spiral artery remodeling is a hallmark of PreE. To better understand EVT interactions with ECs and VSMs, we performed high-resolution re-clustering of these populations and annotated subtypes using canonical markers. This included identification of endovascular EVTs (EVT2), lymphatic ECs (LymphEC), immunomodulatory ECs (ImmEC), and capillary ECs (CapEC) (Fig. 2e–f). Across all EVT subtypes, PreE samples showed a significantly higher proportion of VGLL3⁺ cells compared to HD (Fig. 2g). To assess whether VGLL3 expression influenced cell-cell communication, we conducted CellPhoneDB analysis (32) and observed that VGLL3⁺ EVTs engaged in more ligand-receptor interactions than VGLL3⁻ EVTs (Fig. 2h-i). Notably, these enhanced interactions were concentrated between VGLL3⁺ EVTs and remodeling VSM cells, mature VSMs, and ImmECs (Fig. 2h). Many of the VGLL3⁺-specific interactions were enriched for collagen-related ligands, fibrotic signaling molecules, and components of the WNT signaling pathway (Fig. 2j), supporting a role for VGLL3 in promoting fibrosis and vascular dysfunction in PreE.

To define the gene network centered around VGLL3 in trophoblasts, we analyzed VGLL3-correlated gene expression across 80 trophoblast microarray samples. To stratify 6,634 genes with positive correlation to VGLL3 (Spearman’s ρ > 0), we applied a graphical method identifying a critical inflection point in the correlation decay curve, which yielded a refined set of 737 genes with strong VGLL3 correlation (ρ ≥ 0.67) that constituted a localized transcriptional sub-network (Sup. Fig. 7d, top). Among the top VGLL3-correlated genes were *PSG2*, *PSG3*, *TGFBR2*, and *CSF2RB* (ρ ≥ 0.88). Functional enrichment analysis revealed that this VGLL3-associated network was highly enriched for Gene Ontology biological processes including “granulocyte activation,” “cell activation involved in immune response,” “neutrophil degranulation,” and “myeloid leukocyte-mediated immunity” (Sup. Fig. 7d, bottom). To directly assess the transcriptional impact of VGLL3, we performed bulk RNA-seq following VGLL3 overexpression or knockdown in vitro. Differential expression analysis identified several key PreE-associated genes, *C3*, *AXL*, *CTGF*, and *CYR61*, among the top VGLL3-regulated targets (Fig. 2k), underscoring VGLL3’s role as a potential upstream regulator of pathogenic gene expression in PreE.

### Placental overexpression of Vgll3 is sufficient to trigger PreE-like phenotype in pregnant mice

To assess whether placental overexpression of *Vgll3* is sufficient to induce PreE-like symptoms, we generated a Cre/Lox-based mouse model in which *Vgll3* overexpression was restricted to placental trophoblasts via *Cyp19*-Cre activation during pregnancy (28) (Fig. 3a). Placentas from *Vgll3*-overexpressing (OE) mice exhibited elevated *Flt1* expression, mirroring the molecular phenotype observed in human PreE placentas (Fig. 3b). Pregnant dams carrying *Vgll3*-OE placentas developed gestational hypertension (Fig. 3c), accompanied by reduced cardiac ejection fraction (Sup. Fig. 8a). Histological examination revealed fibrinoid necrosis and microthrombi formation around placental capillaries (Fig. 3d), along with immune cell infiltration (Sup. Fig. 8b). Maternal blood profiling showed elevated levels of mean platelet volume, hematocrit, total protein, blood urea nitrogen, triglycerides, and red blood cell distribution width (Fig. 3e), while placental growth factor (PlGF) was significantly reduced (Sup. Fig. 8c)—a hallmark feature of human PreE. Offspring from *Vgll3*-OE placentas exhibited a 40% postnatal mortality rate (Fig. 3f), reduced body weight at one week of age (Fig. 3g), and lower adiposity by three weeks (Fig. 3h). These findings demonstrate that trophoblast-specific *Vgll3* overexpression is sufficient to induce PreE-like pathology and impair fetal growth.

**Figure 3.**
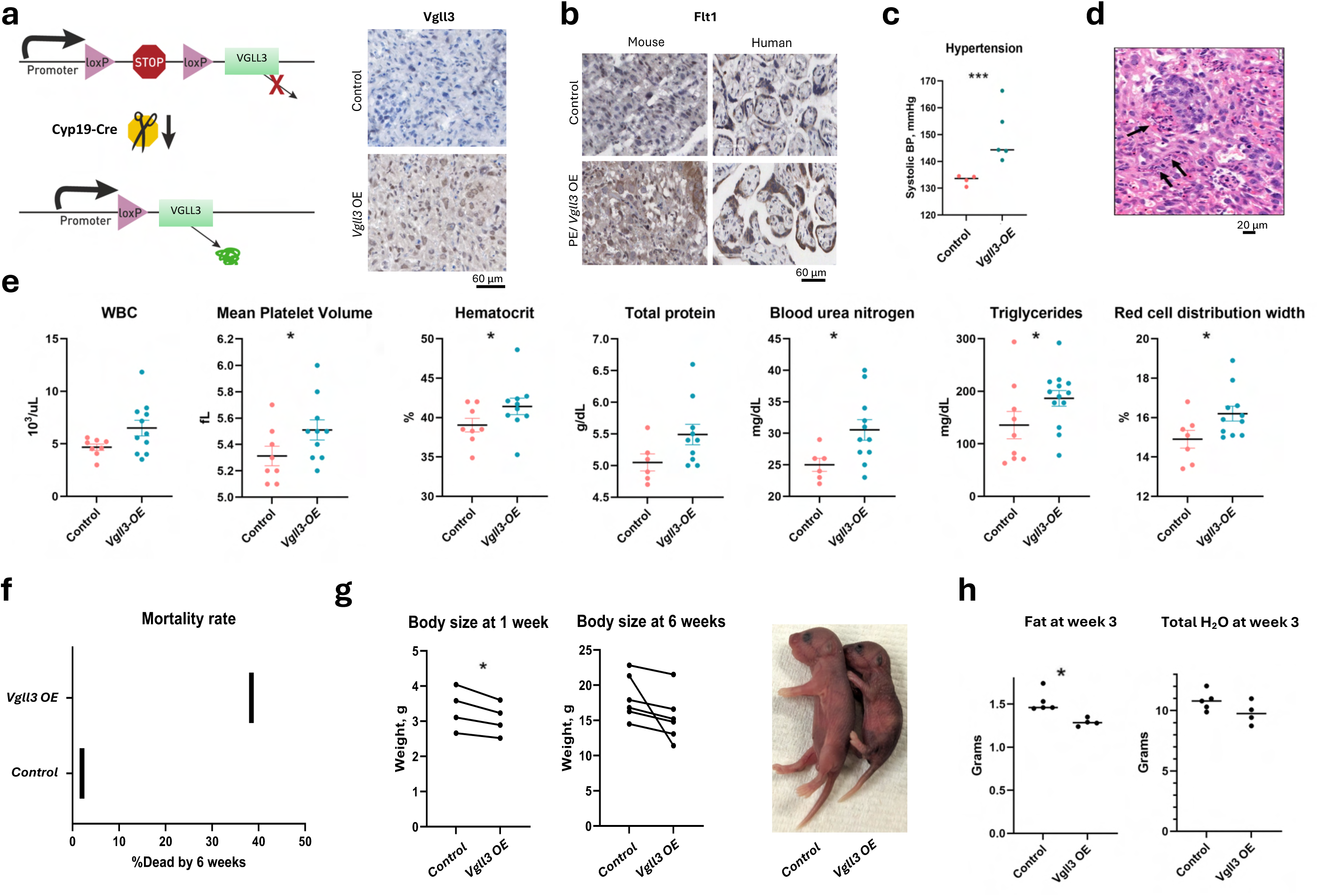
Placental overexpression of *Vgll3* is sufficient to trigger PreE-like phenotype in pregnant mice. a. Strategy for genetically engineering mice with *Vgll3-OE* placentas. Increased levels of Vgll3 were confirmed by IHC (right). b. Upregulation of FLT1 was confirmed both in human PreE placentas and murine *Vgll3-OE* placentas by IHC. c. Elevation in systolic blood pressure (BP) in pregnant dams with Vgll3-OE placentas (*** p < 0.0001) determined by a tail-cuff method at embryonic day 16.5. d. H&E staining revealed microthrombi and fibrinoidnecrosis (arrows) around capillaries in the *Vgll3-OE* placentas. e. Circulatory white blood cells (WBC) and other blood parameters measured in late gestation (* p ≤ 0.05). f. Pups born from *Vgll3-OE* placentas exhibited increased mortality. g. Pups born from *Vgll3-OE* placentas exhibited smaller body size. h. Body composition was measured at 3 weeks (* p ≤ 0.05).

To evaluate the impact of *Vgll3* overexpression on the placental transcriptome, we performed scRNA-seq on control and *Vgll3*-OE mouse placentas (Fig. 4a). Cell clusters were annotated using both mouse-specific marker genes and human signature scores (Fig. 4b). While mouse placenta does not have the same type of cells as human placenta, the trophoblast compartment included trophoblast giant cells (TBGCs) and spongiotrophoblasts (SpongTBs). In *Vgll3*-OE placentas, trophoblasts showed elevated expression of *Flt1*, *Igf2*, and *H19*, mirroring human PreE trophoblasts (Fig. 4c), along with increased activation of the mTOR signaling pathway (Sup. Fig. 9a-b). Immune cells—B cells, myeloid cells, and T cells—also exhibited transcriptional changes consistent with human PreE (Fig. 4d).

**Figure 4.**
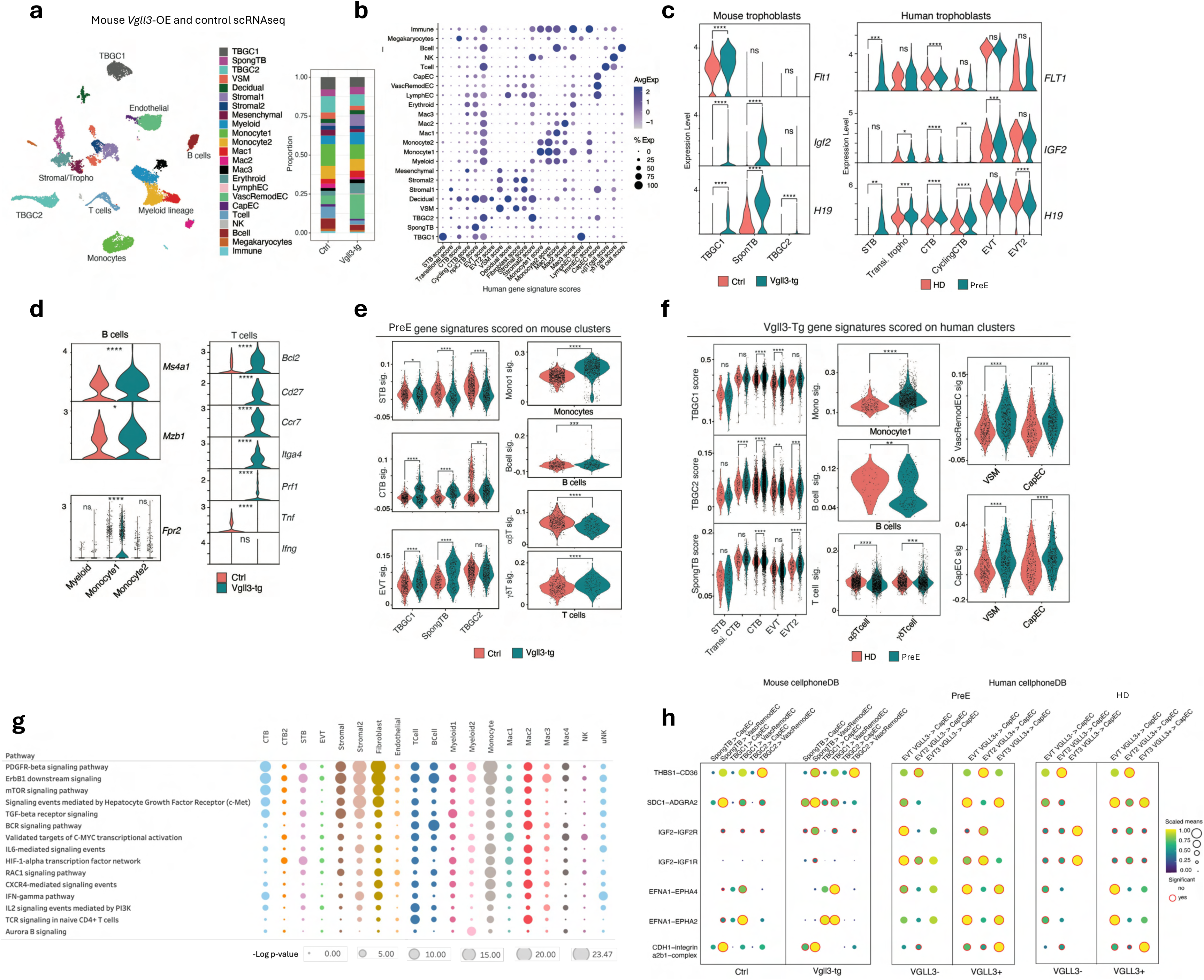
Cells from Vgll3-OE placentas display PreE-associated transcriptomic changes. a. UMAP plot showing mouse placental cells colored by subtypes (left), proportions of clusters in each genotype (right). b. Dot plot of signature scores derived from human clusters in Fig 1a. c. Violin plots showing expression of indicated genes in mouse trophoblasts and their corresponding orthologs in human trophoblasts. (d.-e.) Violin plots showing expression of indicated mouse genes (d.) or module scores (e.) in select clusters separated by genotype. f. Violin plots showing module scores of human ortholog of *Vgll3-OE* specific signatures on select human clusters. g. Double-axis scatter plot of pathways dysregulated in *Vgll3-OE* placentas based on NCI-Nature 2016 database. h. Dot plots show scaled mean scores of indicated interactions between trophoblast lineage cells and endothelial cells in mice (r, left) and human (r, right).

To assess PreE signature enrichment, we identified PreE-specific genes (log₂FC > 0.1, p < 0.05) from human EVT, STB, and cytotrophoblast (CTB) clusters and scored their mouse orthologs across mouse cell types (Fig. 4e, Sup. Table 3). Trophoblasts and monocytes from *Vgll3*-OE placentas displayed significant upregulation of these PreE signatures (Fig. 4f). A reciprocal analysis using a *Vgll3*-OE mouse signature further confirmed elevated scores in human PreE CTB, EVT, and monocyte clusters compared to HD samples (Fig. 4e). These findings indicate that *Vgll3* drives conserved PreE-associated transcriptional programs in both trophoblasts and monocytes across species. Pathway analysis in *Vgll3*-OE placentas identified PDGFR-β, ErbB1, and mTOR signaling as top dysregulated pathways in these cell types (Fig. 4g).

Lastly, we analyzed ligand-receptor interactions in the mouse dataset and found that trophoblast–capillary EC interactions involving receptors critical for vascular integrity and angiogenesis were significantly enriched in dams with *Vgll3*-OE placentas compared to controls (Fig. 4h). These same interactions were also elevated in human VGLL3⁺ EVTs relative to VGLL3⁻ EVTs (Fig. 4h). Together, these findings demonstrate that *Vgll3* overexpression in placentas induces PreE-like phenotypes and provides a viable in vivo model to study the cellular and molecular mechanisms underlying PreE.

### Deletion of Vgll3 suppresses placental inflammation in mice but does not disrupt pregnancy progression

To assess whether VGLL3 is essential for normal pregnancy, we monitored whole-body *Vgll3* knockout (KO) mice throughout gestation. *Vgll3* KO dams exhibited no overt abnormalities, and their pregnancies and offspring were comparable to wild-type (WT) controls. Transcriptomic profiling of KO placentas revealed distinct gene expression clustering (Fig. 5a). Among the most upregulated genes in *Vgll3*-null placentas was *Pcsk6* (Fig. 5b), a gene previously implicated in blood pressure regulation in PreE, with *Pcsk6* deficiency linked to salt-sensitive hypertension (29). Enrichment analysis of upregulated genes pointed to pathways involved in “positive regulation of cell migration” and “regulation of angiogenesis,” whereas downregulated genes were associated with “signaling events mediated by VEGFR1 and VEGFR2” (Fig. 5c).

**Figure 5.**
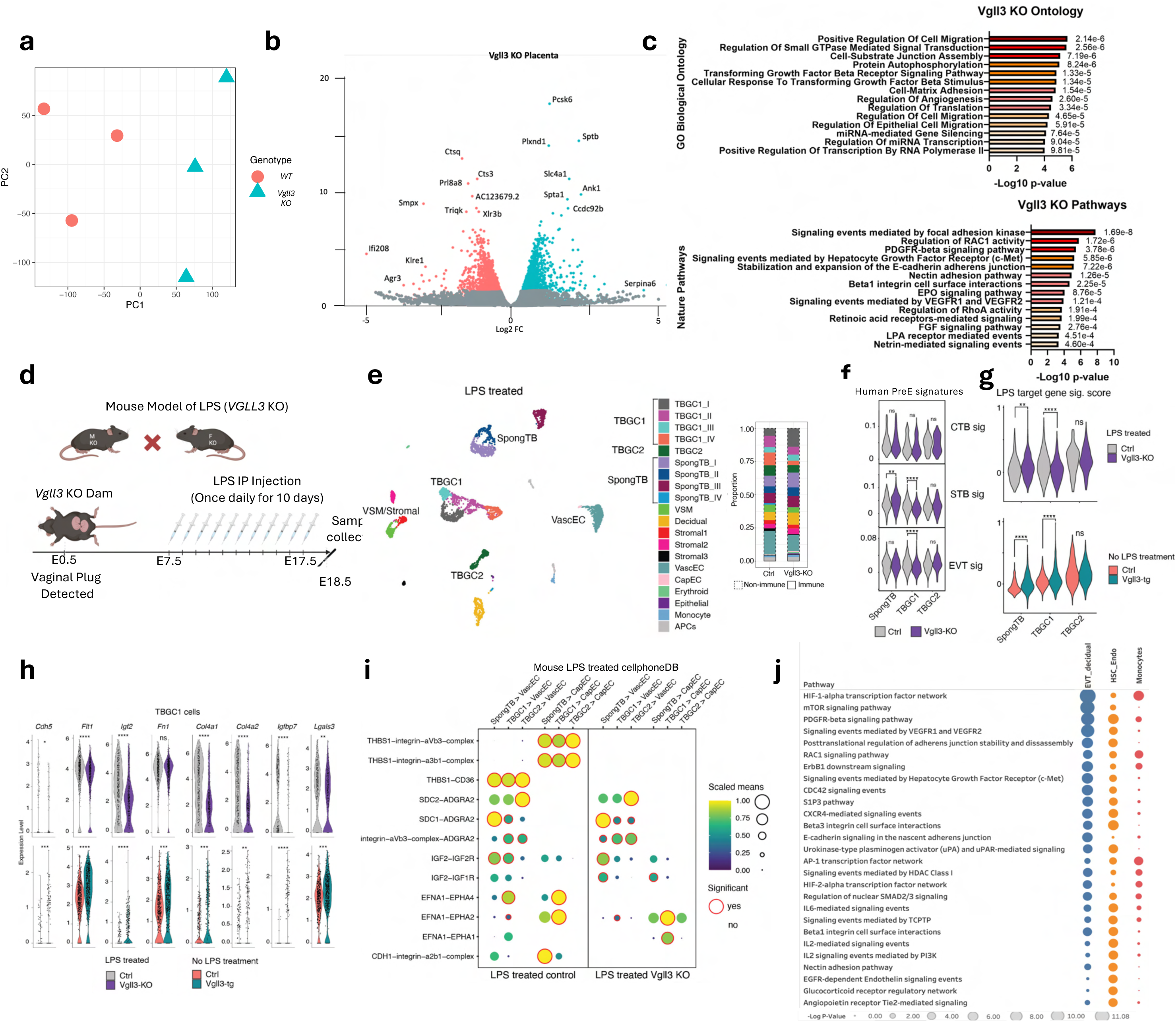
Deletion of *Vgll3* suppresses placental inflammation in mice. a. Principal component (PC) scatter plot shows distinct separation of placenta transcriptomes from murine *WT* and *Vgll3 KO* mice. b. Volcano plot with top DEGs in *Vgll3 KO* placentas (n=3 per group). c. Bar graphs with top dysregulated pathways in *Vgll3 KO* placentas per GO Biological Process 2023 and NCI-Nature 2016 databases. d. Mouse model of placental inflammation in the absence of *Vgll3*. e. UMAP plot showing cells from placenta of LPS treated *WT* and *Vgll3 KO* mice and their proportions. f. Human PreE-specific gene expression module within major trophoblast populations scored across murine trophoblasts from *WT* or *Vgll3 KO* mice treated with LPS. g. Module score of LPS-targeted genes was calculated in murine placenta trophoblasts from either (top) *WT* or *Vgll3 KO* mice treated with LPS or (bottom) untreated *WT* or *Vgll3-OE* mice. h. Violin plots showing expression of indicated PreE-associated genes in TBGC1 trophoblasts. i. Dot plots show scaled mean scores of indicated interactions between trophoblast lineage cells and endothelial cells in LPS treated mice. j. Double-axis scatter plot of pathways differentially regulated in placentas from LPS-administered *Vgll3 KO* mice based on NCI-Nature 2016 database.

Our analyses identified lipopolysaccharide (LPS) as a potential upstream regulator of the PreE-specific transcriptomic shift in human placentas (Fig. 1b), consistent with previous studies in which LPS was used to induce PreE-like inflammation in animal models (30). To assess whether *Vgll3* deletion mitigates LPS-induced inflammation, we administered daily LPS injections (Sigma-Aldrich L2630, 20 μg/kg) to pregnant WT and *Vgll3* KO dams from embryonic day 7.5 to 17.5 (Fig. 5d) and performed scRNA-seq on placentas collected at day 18.5 (Fig. 5e, left). Deletion of *Vgll3* did not substantially alter the composition of major placental cell clusters (Fig. 5e, right). However, scoring with a human PreE-specific gene signature revealed that TBGC1 cells from *Vgll3* KO mice exhibited reduced expression of both STB and EVT PreE gene signatures (Fig. 5f). Moreover, *Vgll3* KO trophoblasts showed decreased expression of an LPS-induced gene signature (Fig. 5g, top). In contrast, trophoblasts from *Vgll3*-OE placentas (in the absence of LPS) displayed increased expression of this same LPS signature compared to controls (Fig. 5g, bottom). These transcriptional changes were accompanied by a modest, though not statistically significant, reduction in blood pressure in LPS-treated *Vgll3* KO dams compared to LPS-treated WT controls (Sup. Fig. 10).

Further supporting the role of *Vgll3* in regulating *Flt1* and cell adhesion molecule expression in trophoblasts, we observed reduced expression of these genes in trophoblasts from LPS-injected *Vgll3* KO dams compared to LPS-injected WT controls, and conversely, increased expression in *Vgll3*-overexpressing (OE) placentas compared to WT controls (Fig. 5h). Ligand-receptor interaction analysis revealed that PreE-specific trophoblast–EC interactions identified in the *Vgll3*-OE mouse model were significantly enriched in LPS-treated WT mice but diminished in LPS-treated *Vgll3* KO mice (Fig. 5i). Among the top differentially regulated pathways in LPS-treated *Vgll3* KO placentas were HIF-1α and AP-1 signaling, particularly within EVTs, ECs, and monocytes (Fig. 5j). These findings underscore the importance of *Vgll3* in driving inflammatory and adhesive signaling in trophoblasts and its broader impact on placental cell-cell communication. Given the effectiveness of *Vgll3* deletion in attenuating placental inflammation in mice, we next sought to target VGLL3/Hippo signaling in human PreE placentas.

### Targeting Hippo pathway suppresses disease signaling in PreE placentas ex vivo

While a specific VGLL3 inhibitor is not yet available, verteporfin is a known inhibitor of the Hippo pathway at the transcriptional level (31). To evaluate whether verteporfin mimics the effects of VGLL3 inhibition, we treated HTR8 trophoblasts with either verteporfin or VGLL3-targeting siRNA and performed bulk RNA-seq. Both treatments impacted overlapping pathways, including integrin-mediated cell surface interactions, angiogenesis, p53 effector signaling, and Syndecan-1 signaling (Sup. Fig. 11a).

Comparison of DEGs revealed substantial overlap: 1,191 of the 1,493 DEGs identified in VGLL3 siRNA-treated cells (80%) were also differentially expressed in verteporfin-treated cells (Sup. Fig. 11b). Notably, verteporfin induced broader transcriptomic changes than VGLL3 siRNA, likely reflecting additional VGLL3-independent effects. We categorized DEGs into those shared between treatments and those unique to each condition (Sup. Fig. 11c) and performed pathway enrichment analyses for each group (Sup. Fig. 11d). The overlapping DEGs were enriched in pathways associated with apoptosis, TNF signaling, hypoxia, and mTORC signaling, all previously implicated in human PreE and the Vgll3-OE mouse model.

PreE placentas were treated ex vivo with the Hippo pathway inhibitor verteporfin, followed by single-cell RNA sequencing (scRNA-seq) analysis (Fig. 6a, left). Compared to vehicle (DMSO) controls, verteporfin treatment did not induce major shifts in overall cellular composition (Fig. 6a, right). Receptor-ligand expression analysis demonstrated that verteporfin broadly suppressed PreE-associated intercellular communication, most notably among decidual cells, stromal cells, ECs, and trophoblasts (Fig. 6b). Correspondingly, cytokine signaling was reduced across all major cell types following verteporfin treatment (Fig. 6c). Verteporfin treatment also attenuated PreE gene signature scores, previously derived from comparisons between untreated PreE and HD placentas, in both CTBs and EVTs (Fig. 6d). To further dissect verteporfin’s transcriptional effects, we compared DEGs from disease-state (PreE vs. HD) and treatment (verteporfin vs. DMSO) contrasts in trophoblasts (Fig. 6e). Multiple genes elevated in PreE were downregulated by verteporfin, including IGFBP7, COL17A1, EFNB1, and SEMA4C (Fig. 6e–f). In stromal cells, LTBP4, a regulator of TGF-β activity and a recently identified VGLL3 target in salmon sexual maturation (32), was significantly elevated in PreE compared to HD and downregulated following verteporfin treatment (Fig. 6g).

**Figure 6.**
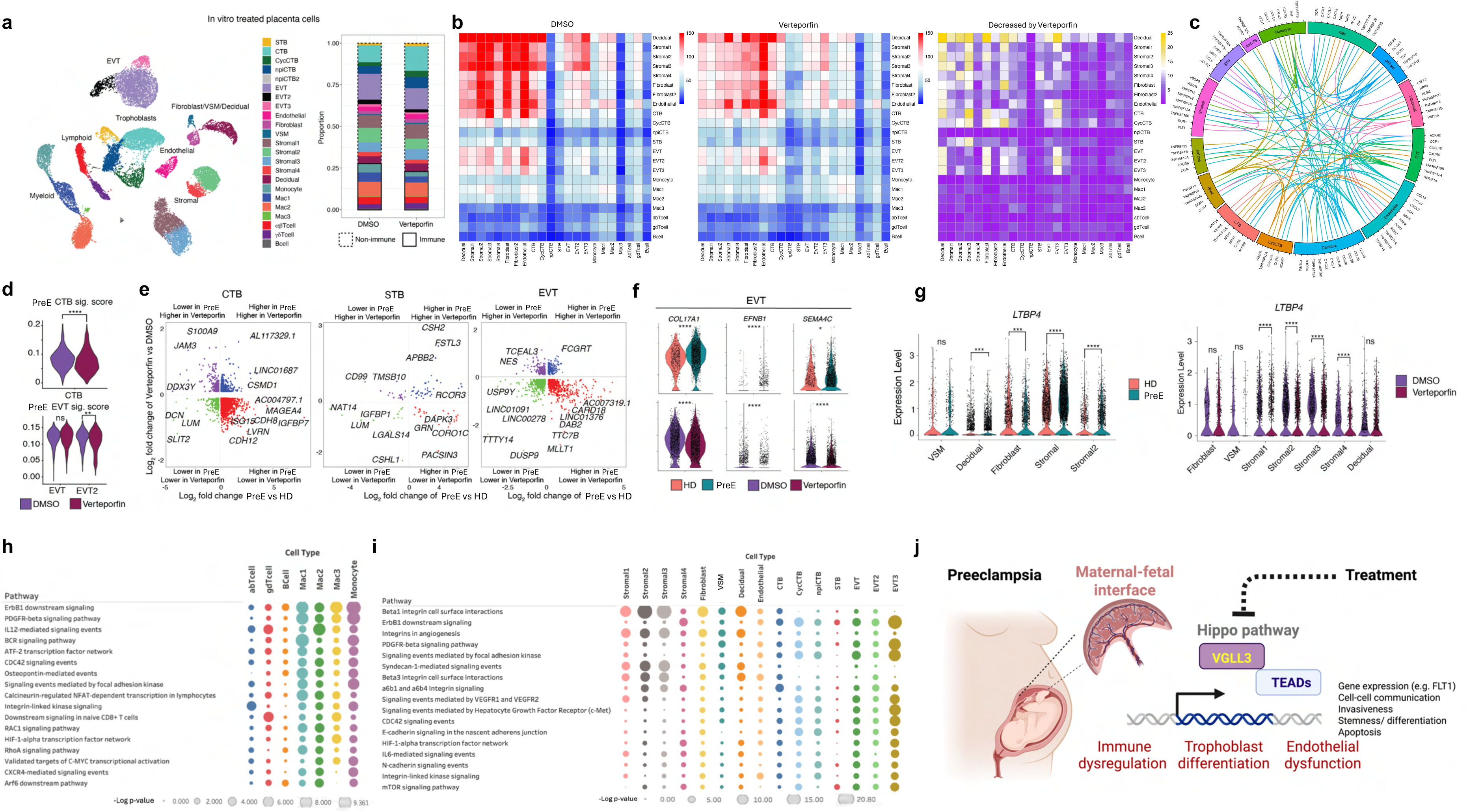
Targeting Hippo pathway suppresses PreE disease signaling *ex vivo*. a. UMAP plot showing human placental cells in placentas obtained from PreE patients and treated with verteporfin or DMSO ex vivo, colored by cluster (left), with bar graph showing proportions of clusters in each treatment (right). b. Heatmaps showing number of ligand-receptor pairs between indicated populations in the two treatment groups (left and center) and as reduced by verteporfin (right). c. Circos plot depicting cytokine interactions reduced by Verteporfin. d. Violin plots show module score of PreE CTB (top) or EVT (bottom) gene signatures on indicated cell clusters in DMSO- or verteporfin-treated placentas. e. Scatter plots show PreE/ HD differential gene expression in indicated CTB, STB, or EVT from untreated samples shown in Fig1 (log2 fold change, x-axis) vs. Verteporfin/DMSO differential gene expression (log2 fold change, y-axis). Red dots indicate cells that are higher in PreE trophoblasts and repressed in verteporfin-treated cells. f. Violin plots show expression of indicated genes in EVT from PreE and HD untreated placenta (top) and in *ex vivo* treated PreE placenta (bottom). g. Violin plots of LTBP4, a gene targeted by VGLL3, in PreE placentas (vs HD controls) and in Verteporfin-treated PreE placentas (vs DMSO-treated controls). h-i. Double-axis scatter plot of pathways differentially regulated by Verteporfin in PreE placentas in immune (l) and non-immune (m) cells as based on NCI-Nature 2016 database. k. Schematic of VGLL3-driven dysregulation at the maternal-fetal interface in PreE.

Pathway enrichment of verteporfin-affected genes in immune cells highlighted suppression of ERBB1, IL-12, and BCR signaling (Fig. 6h), while in non-immune cells, verteporfin perturbed pathways involving integrin-mediated cell-surface interactions, angiogenesis, and FAK signaling (Fig. 6i). Notably, verteporfin-treated placental explants exhibited a substantial overlap in DEGs and pathways with those affected by WRW4, an FPR2 antagonist (Sup. Fig. 12a–b), consistent with prior findings that suppression of FPR2 mitigates inflammation in PreE placentas (33).

Taken together, these findings suggest that the dysregulation of the Hippo pathway in PreE leads to an immune imbalance at the maternal-fetal interface, alters trophoblast differentiation, and causes vascular endothelial dysfunction (Fig. 6j). As these key pathological processes of PreE converge on a VGLL3-centered gene network, our data position VGLL3 as a pivotal master regulator in the etiology of PreE.

## DISCUSSION

It has been hypothesized that the female immune system evolved specialized adaptations to tolerate the immunologically invasive placenta while maintaining heightened immune surveillance against pathogens. While this balance may confer protection against infections and cancer, it also introduces female-specific immune signaling that increases susceptibility to autoimmune diseases. In the context of pregnancy, this immune complexity may contribute to dysregulated maternal-fetal immune interactions, thereby predisposing some women to PreE (34).

Pregnancy represents a unique immunological state in which trophoblast invasion and fetal development depend on a finely tuned local inflammatory environment that supports cell clearance, angiogenesis, cell growth, and immune tolerance (35), processes that are disrupted in PreE (2). The Hippo signaling pathway, an evolutionarily conserved regulator of cell proliferation, differentiation, apoptosis, and mechanotransduction (36), has been implicated in fibrosis and endothelial dysfunction in autoimmune diseases (37). In the context of pregnancy, emerging evidence suggests that the Hippo pathway regulates trophoblast stemness (26, 38), angiogenesis (39), and maternal-fetal immune tolerance (40), and may contribute to pregnancy complications (41), including PreE (42, 43). Through detailed transcriptomic analyses of immune and non-immune cells in PreE placentas, we identify dysregulation of the Hippo pathway as a central feature of disease pathogenesis, with the transcriptional co-regulator VGLL3 emerging as a key mediator linking Hippo signaling to immune activation, trophoblast dysfunction, and endothelial pathology in PreE.

Our group previously identified VGLL3 as a regulator of a pro-inflammatory gene network in the skin that predisposes women to autoimmune diseases (21), including cutaneous lupus and Sjögren’s syndrome, both of which show VGLL3 upregulation. Notably, epidermal overexpression of *Vgll3* in mice is sufficient to induce a lupus-like autoimmune phenotype, characterized by hallmark features such as autoantibody production (20). Beyond autoimmunity, VGLL3 has emerged as a key regulator at the intersection of sex-specific biology and development. Independent studies have linked VGLL3 to the timing of sexual maturity in Atlantic salmon (44) and to age of menarche in humans (45), an important clinical connection, as early menarche is associated with increased risk of both PreE (8, 9) and autoimmune disease(46).

Strikingly, VGLL3 expression is highest in the placenta compared to all other organs. Although VGLL3 lacks a DNA-binding domain, it exerts its transcriptional effects by binding TEAD transcription factors, thereby activating Hippo pathway signaling, a mechanism best characterized in cancer (47, 48). During pregnancy, trophoblast proliferation, differentiation, migration, and invasion of the maternal uterus and vasculature mirror features of malignant tumors, a phenomenon termed “trophoblast pseudo-tumorigenesis” (49). Our data suggest that VGLL3 may play a physiological role in placental development through its regulation of these processes. However, in PreE, VGLL3 appears to be dysregulated: its overexpression and interaction with TEAD1 and TEAD3 drive immune dysregulation and trophoblast dysfunction, ultimately contributing to vascular pathology and pregnancy-induced hypertension.

In summary, our findings reveal that VGLL3 acts as a shared molecular nexus linking PE, sexual maturation, and autoimmunity. This provides a mechanistic explanation for the long-recognized clinical associations between PE, early menarche, and autoimmune disorders. Our study does have limitations, including the current lack of data defining the upstream drivers of VGLL3 dysregulation in PE. Preliminary data from our lab suggest a role for epigenetic regulation via X-chromosome–encoded histone demethylases (unpublished). Additionally, the individual contributions of TEAD1 and TEAD3 in PE remain to be elucidated, offering compelling directions for future research. Nevertheless, our findings position VGLL3 and its associated signaling network as a promising therapeutic target in the treatment of PE.

## RESOURCE AVAILABILITY

### Lead Contact

Requests for further information and resources should be directed to and will be fulfilled by the lead contact, Johann E. Gudjonsson.

### Materials Availability

Mouse lines and any other unique/ stable reagents generated in this study will be made available on request.

### Data and Code Availability

Sequencing data have been deposited at NCBI Gene Expression Omnibus as GSE298602 and are publicly available as of the date of publication.

## ACKNOWLEDGEMENTS

We would like to thank Drs. Guðrún Valdimarsdóttir, Nándor Gábor Than, and Smarajit Mondal for their helpful scientific discussions, Drs. Enze Xing, Xianying Xing, Vincent Van Drongelen, Mehrnaz Gharaee-Kermani, Ranjitha Uppala, and Nitin Kumar for their assistance, Dr. Gustavo Leone for generously sharing Cyp19-Cre mice (28), Steven Whitesall for help with BP monitoring and Jennifer Fox for help with sequencing submissions. We would also like to acknowledge University of Michigan Advanced Genomics Core, Epigenomics Core, In-Vivo Animal Core, Proteomics Resource Facility, Physiology Phenotyping Core, Animal Metabolic, Physiological Behavioral Phenotyping Core, Unit for Laboratory Animal Medicine, Skin Research Center, and Frankel Cardiovascular Center’s Michigan Biological Research Initiative on Sex Differences in Cardiovascular Disease (M-BRISC). This work was supported by the Taubman Medical Research Institute, NIH-P30-AR075043 (JEG, AAD, MK, LCT), NIH-R01-AI183620 (JEG), K24AR076975 (JMK), K08-AR078251 (ACB), and M-BRISC pilot grant (OP).

## AUTHOR CONTRIBUTIONS

O.P. and J.E.G. conceived experiments, wrote the manuscript, and secured funding. O.P., R.P., K.Y., L-J.S., J.E., J.K., L.B.C., H.Z., R.B., W.R.S., L.C.T. performed experiments, including data analyses. A.S., E.S.L. provided human samples. I.L.B., M.K.S., N. L. W., K. S., K.A.G., J.M.K., A.C.B., A.A.D., S.K.G., L.C.T., J.E.G. provided expertise and feedback.

## SUPPLEMENTAL FIGURE TITLES AND LEGENDS

**Supplemental Figure 1.**
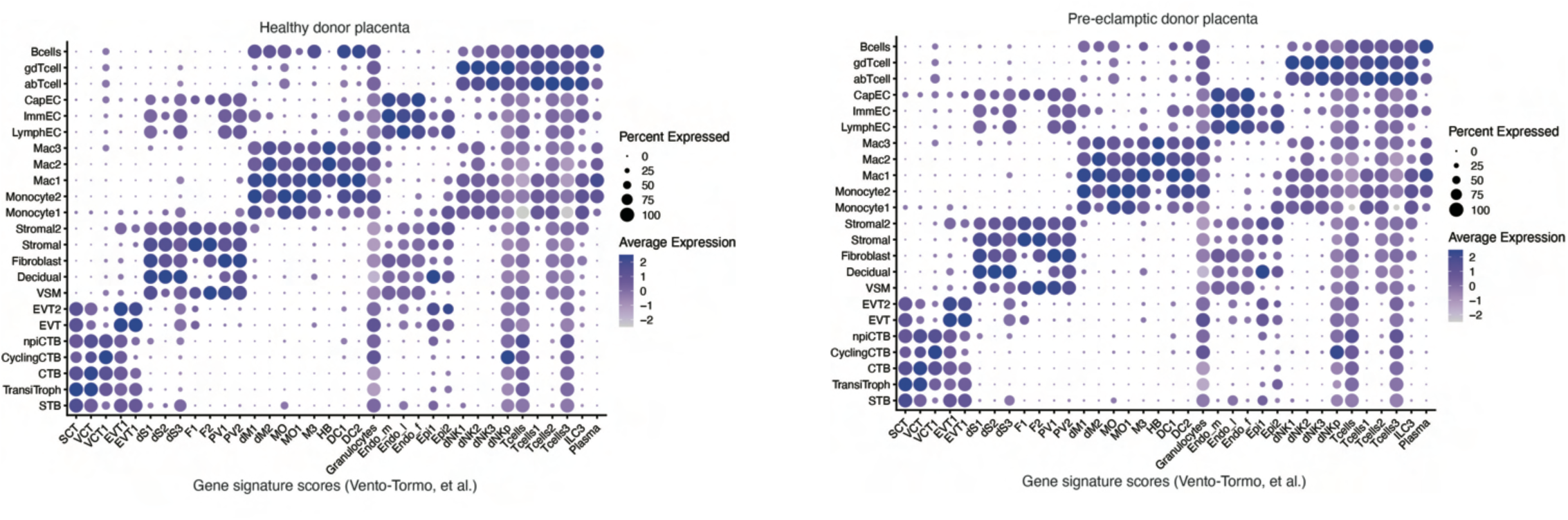
Gene signature scores for cell type assignments in human scRNA-seq data.

**Supplemental Figure 2.**
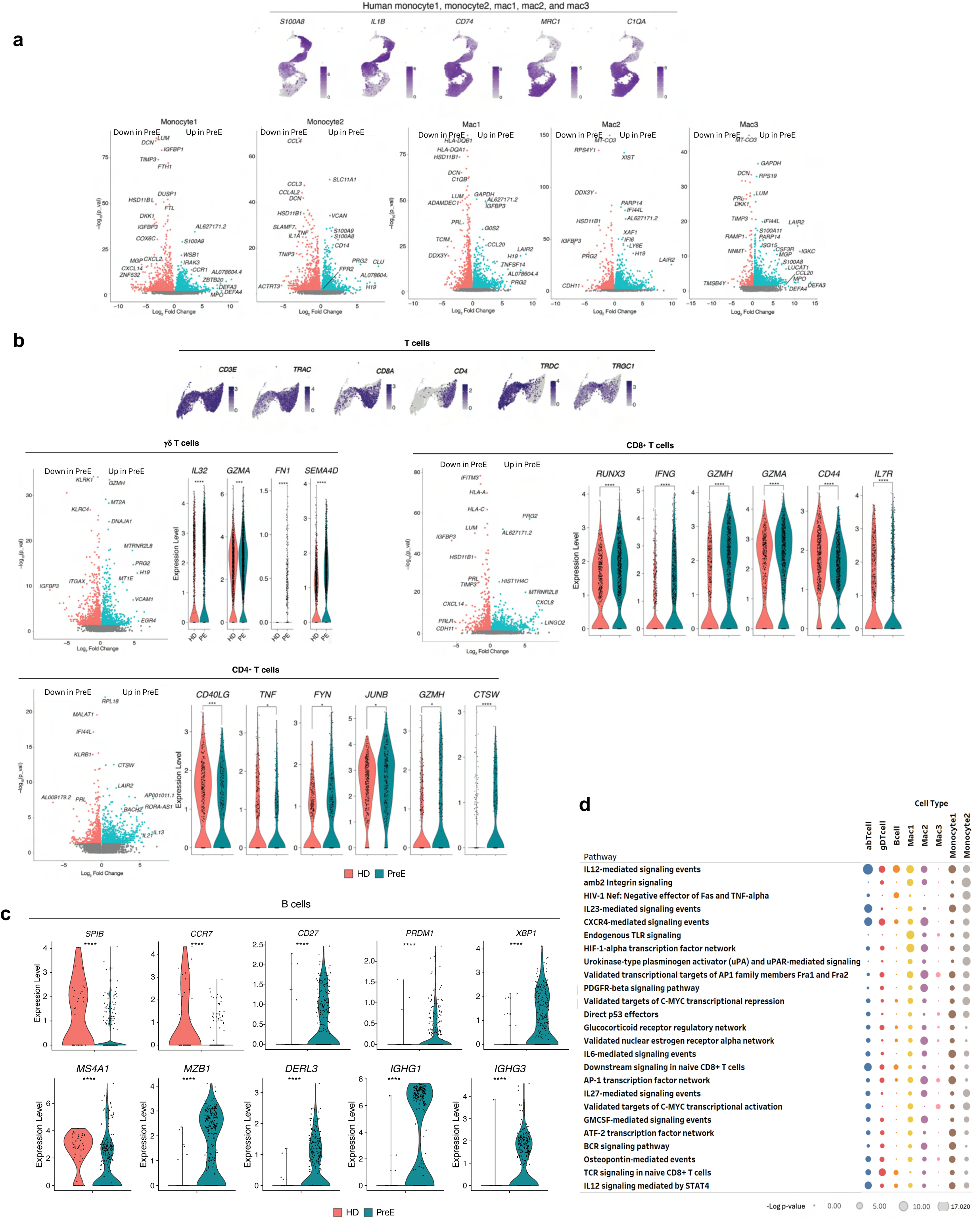
a. Scaled expression of indicated genes is shown on UMAP plots of myeloid lineage cells. Volcano plots show differential gene expression (Log2 fold-change [x-axis] vs. P value, - Log10 scale [y-axis]) between HD and PreE in indicated myeloid clusters. b. Markers for identifying major T cell sub-types (top). Volcano plots and violin plots of key genes (bottom) in γδ T cells (left), CD8+ T cells (middle), and CD4+ T cells (right). c. Violin plots of select DEGs in PreE B cells. d. Double-axis scatter plot of pathways dysregulated in PreE placental immune cells based on NCI-Nature 2016 database.

**Supplemental Figure 3.**
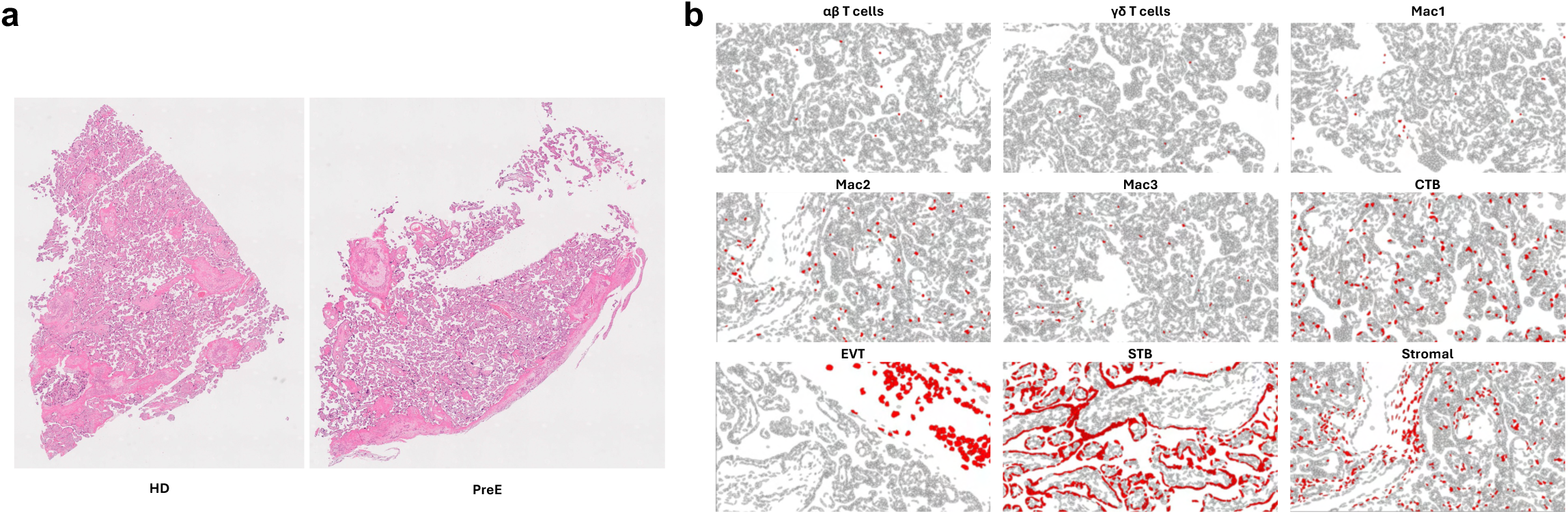
a. H&E staining of HD and PreE placenta samples used for spatial seq. b. scRNA-seq cell clusters overlayed onto Xenium spatial seq data. PreE placenta villi are shown.

**Supplemental Figure 4.**
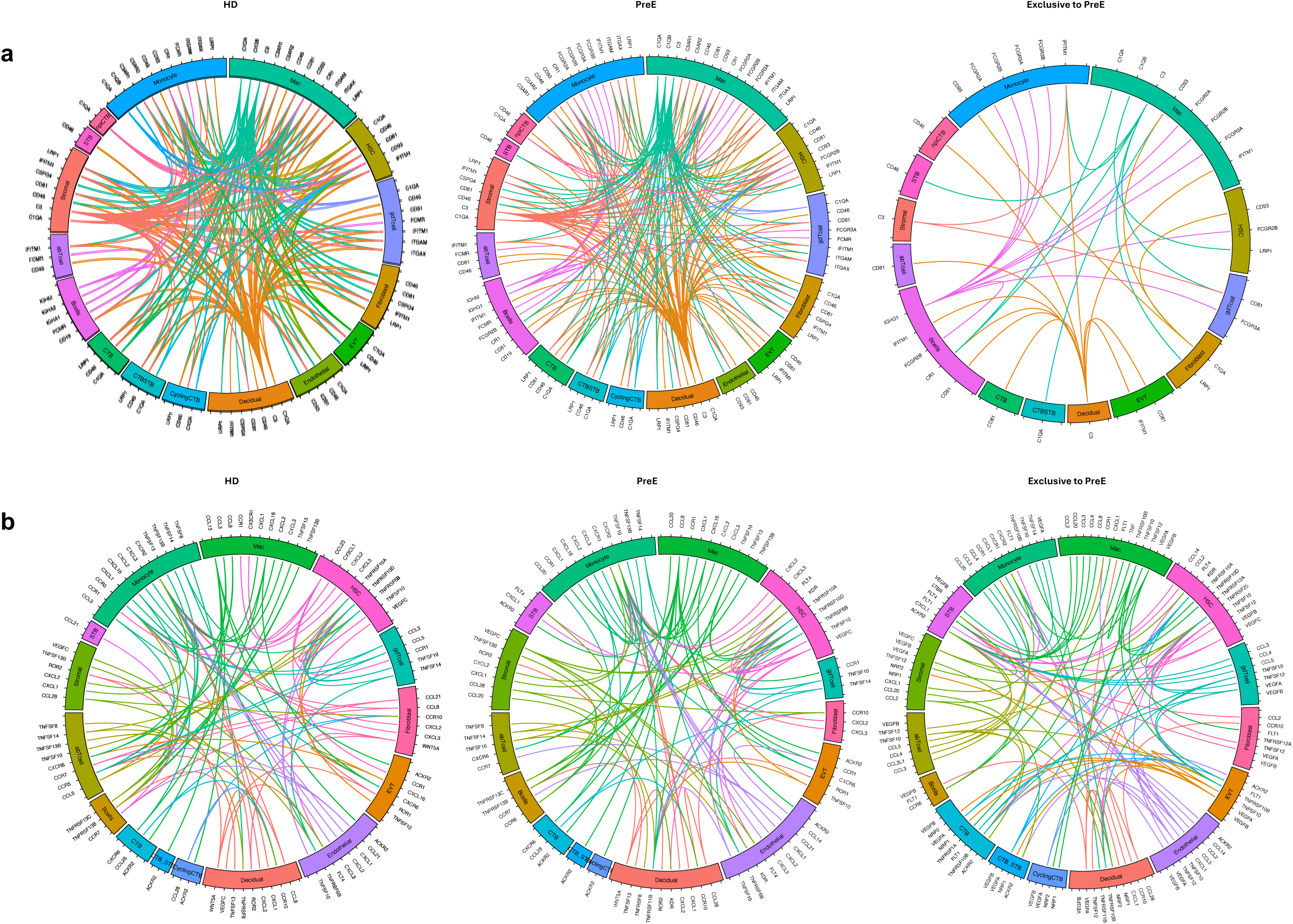
a. Circos plots depicting PreE-specific complement/ immunoglobulin interactions in major cell types. b. Circos plots depicting PreE-specific cytokine interactions in major cell types.

**Supplemental Figure 5.**
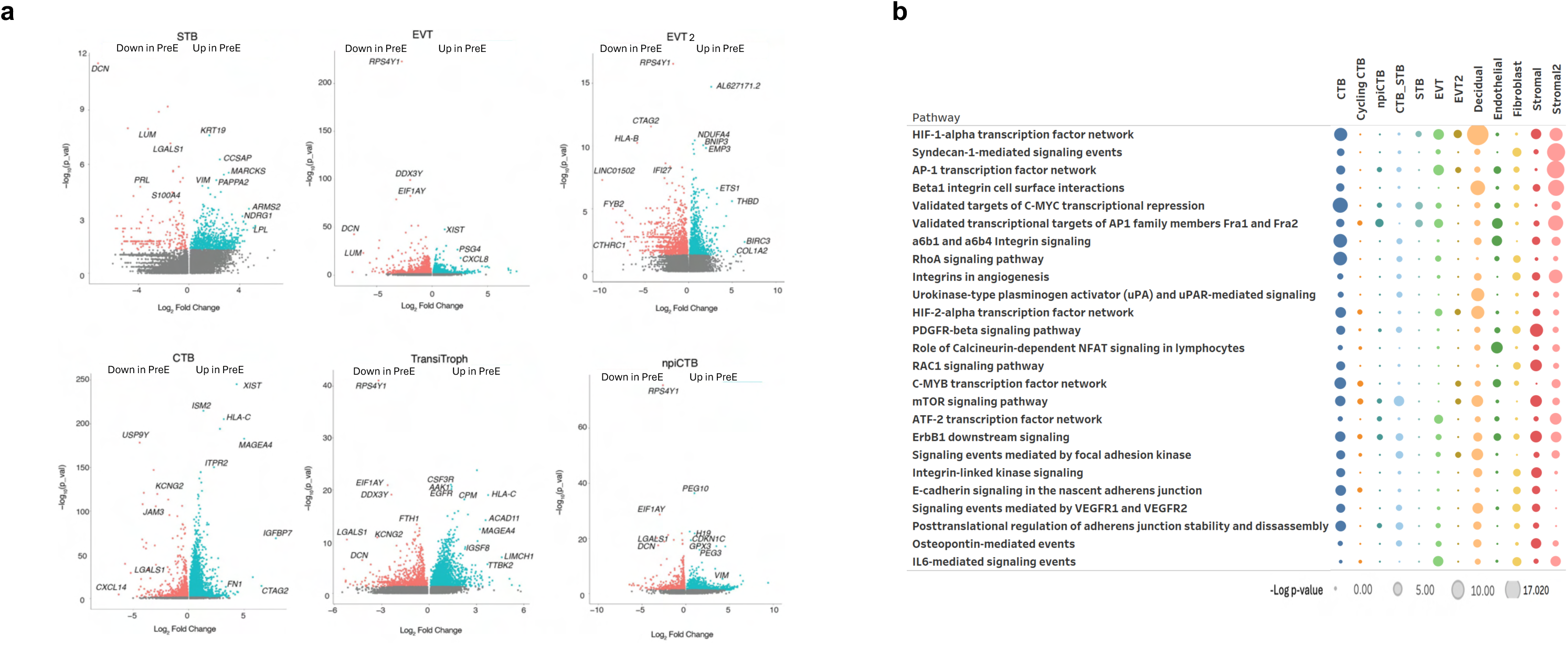
a. Volcano plots of DEGs in main trophoblast types of PreE placentas vs HD. b. Double-axis scatter plot of pathways dysregulated in PreE placental non-immune cells based on NCI-Nature 2016 database.

**Supplemental Figure 6.**
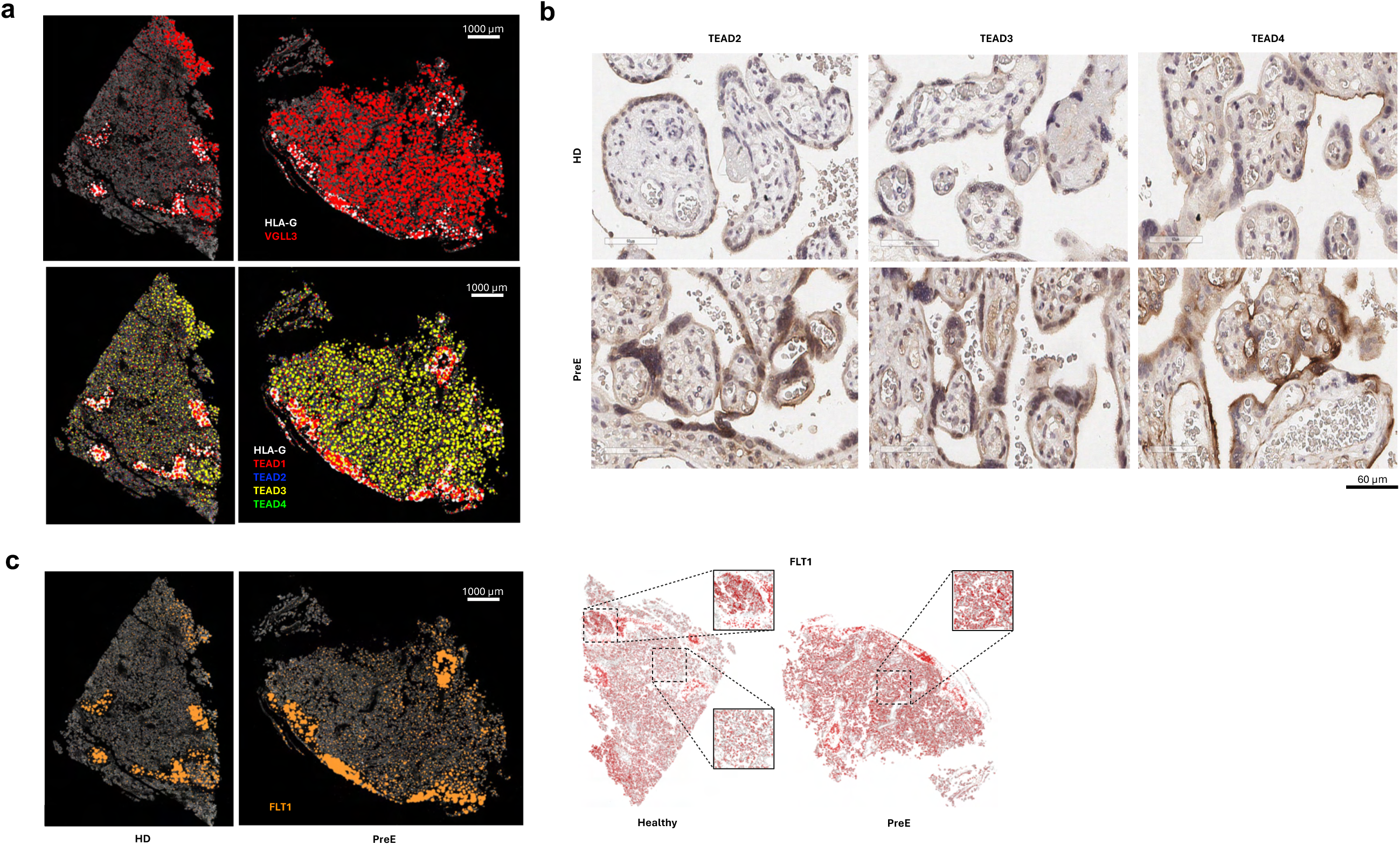
a. Expression patterns of VGLL3, and TEADs along with a decidual marker HLA-G as determined by spatial seq. b. IHC micrographs showing upregulated TEAD2-4 expression in PreE placentas. c. A whole-tissue and zoomed-in visualization of FLT1 expression.

**Supplemental Figure 7.**
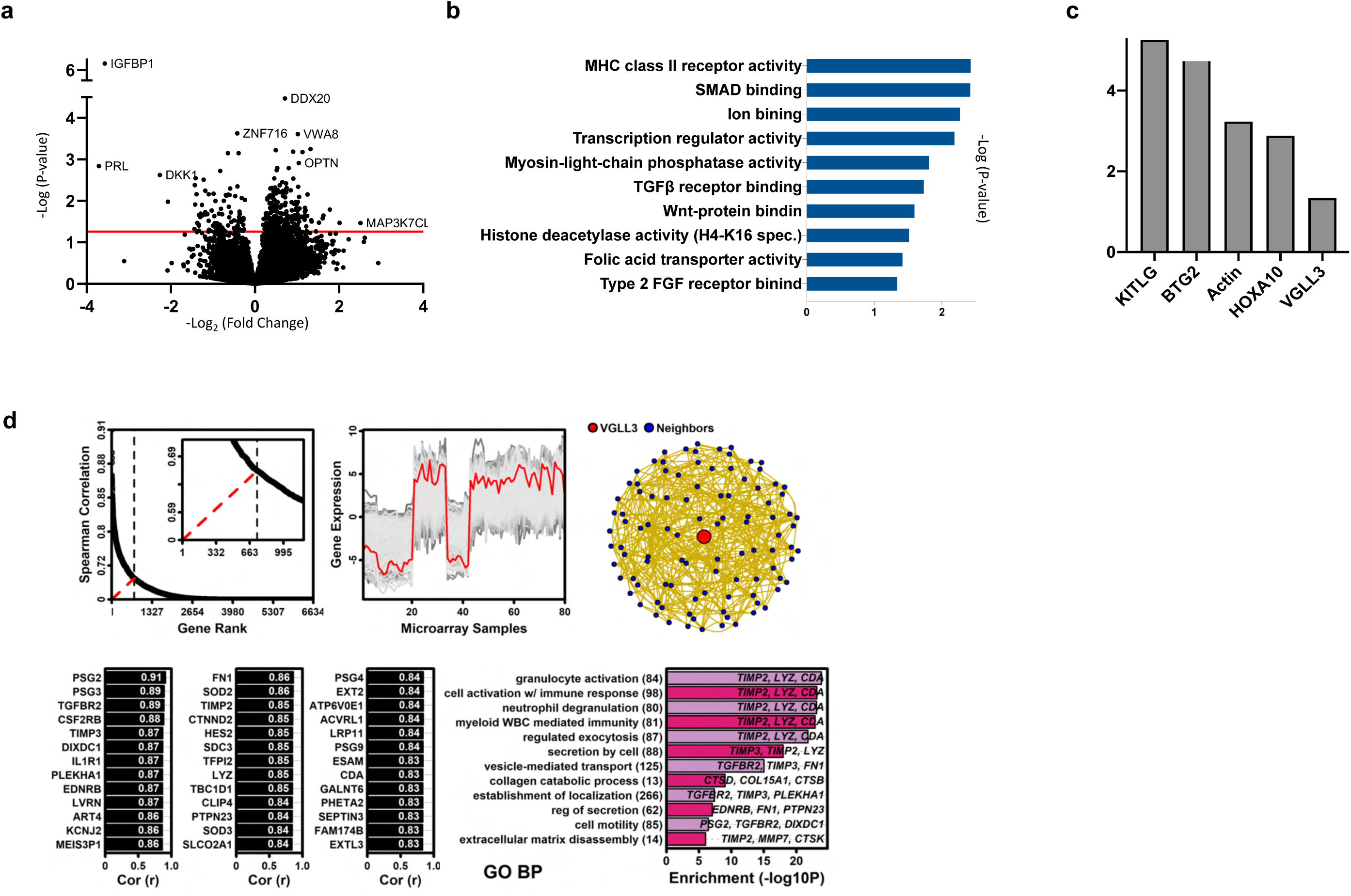
a. Volcano plot with top DEGs in FFPE PreE placentas (n=43) vs matched HD controls (n=41). b. Top pathways from 259 up- and 90 down-regulated genes with statistical significance of p ≤ 0.05 in a. c. Select upstream regulators identified by IPA. d. VGLL3-correlated identified from 80 trophoblast microarray samples. Out of 13,753 genes with detectable expression (p < 0.05, Signed rank test) in at least 5% samples, the expression pattern of 6634 was positively correlated with VGLL3 (rs > 0). Spearman rank correlation estimates varied continuously among these genes. To determine an appropriate threshold, a graphical approach was used that identified a critical point in the decay of correlations and defined a set of 737 genes with VGLL3-correlated expression (rs ≥ 0.67), representing a local sub-network surrounding VGLL3. Genes most strongly correlated with VGLL3 and pathways they are enriched in are listed. The number of genes associated with each term is shown in parentheses.

**Supplemental Figure 8.**
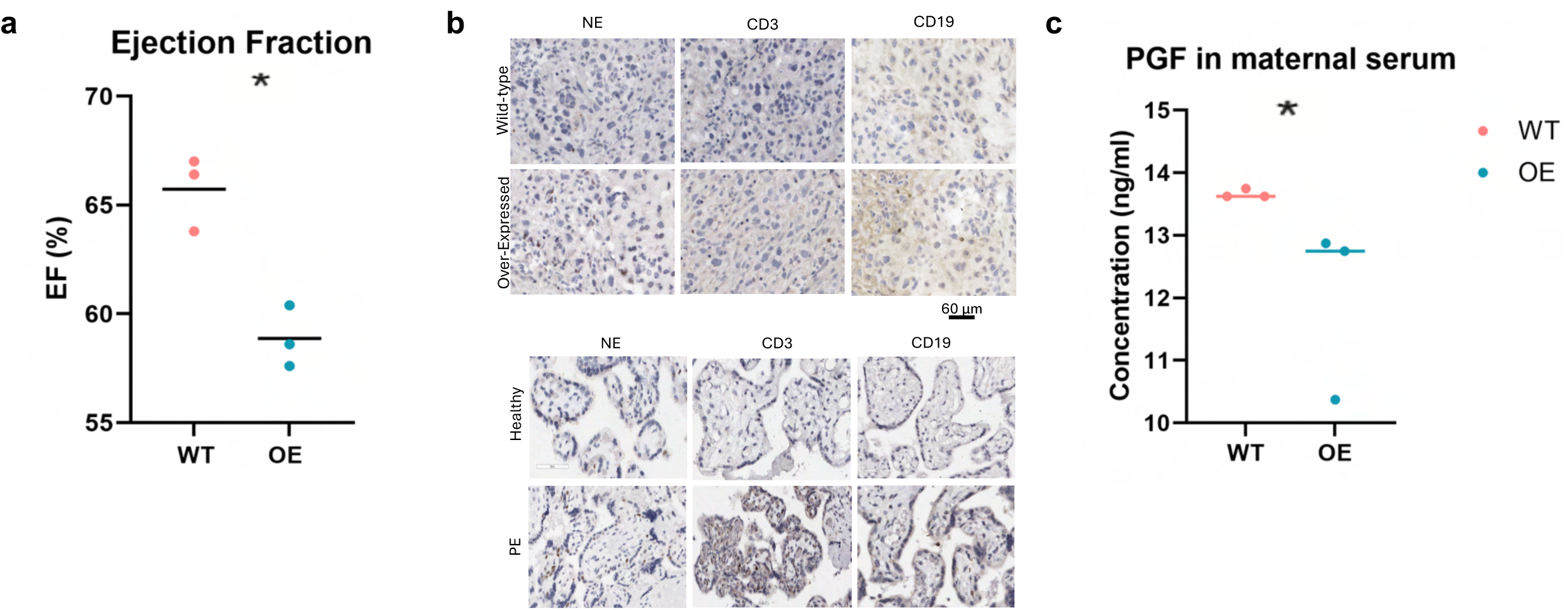
a. Ejection fraction measured at E15.5 using ultrasound with Echo. b. IHC detected neutrophils, T cells, and B cells in *Vgll3-OE* placentas (top) and corresponding cell populations in human PreE placentas (bottom). c. PGF levels in pregnant dams with *Vgll3-OE* placentas as determined by ELISA.

**Supplemental Figure 9.**
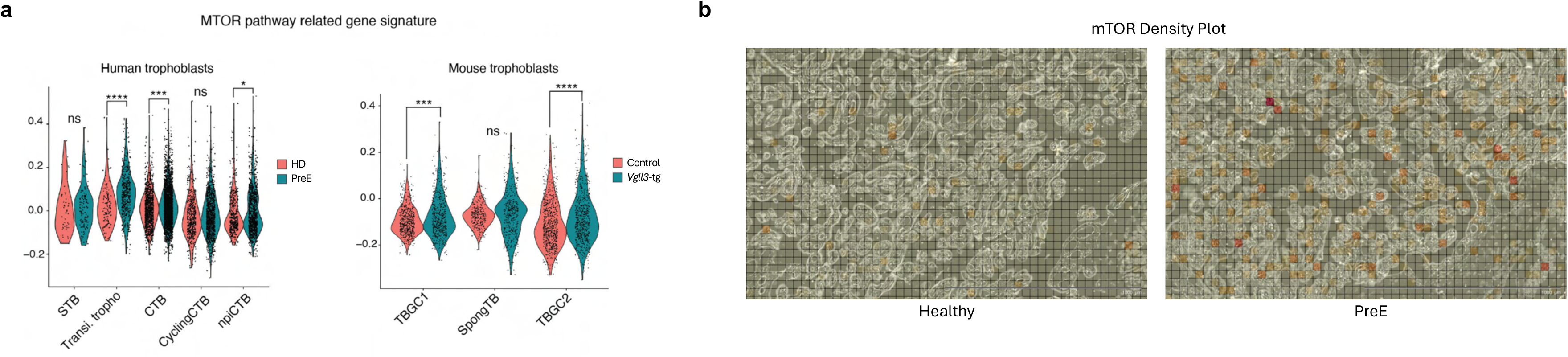
a. MTOR gene signature is increased in trophoblasts from human PreE placentas and mouse *Vgll3-OE* placentas. b. Spatial seq illustrates upregulation of mTOR in PreE placenta.

**Supplemental Figure 10.**
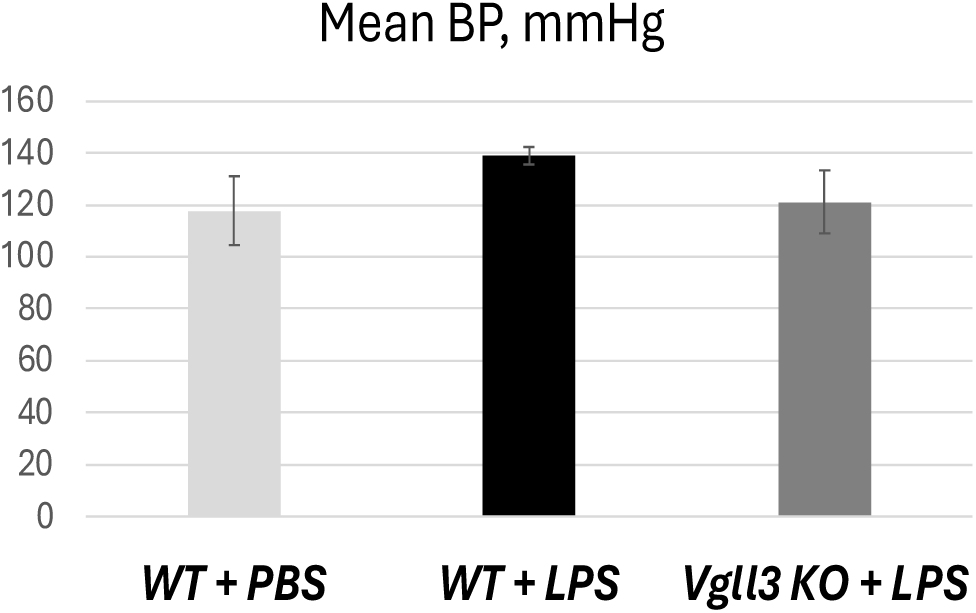
Mean BP for LPS-treated mice and PBS-treated controls.

**Supplemental Figure 11.**
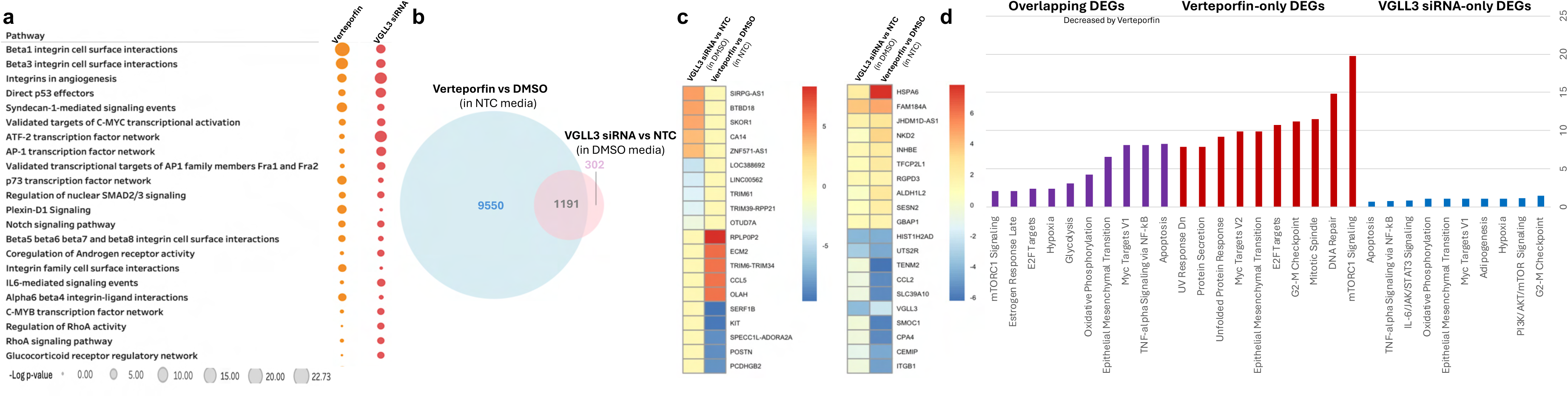
a. Double-axis scatter plot of pathways dysregulated in HTR8 trophoblasts treated with either Verteporfin (vs DMSO) or VGLL3 siRNA (vs NTC), based on total DEGs imputed into NCI-Nature 2016 database. b. Number of unique and overlapping DEGs (p < 0.05) for two treatment groups from a. c. Top unique and overlapping DEGs are listed. d. Top pathways for Verteporfin-only, VGLL3 siRNA-only, and overlapping DEGs.

**Supplemental Figure 12.**
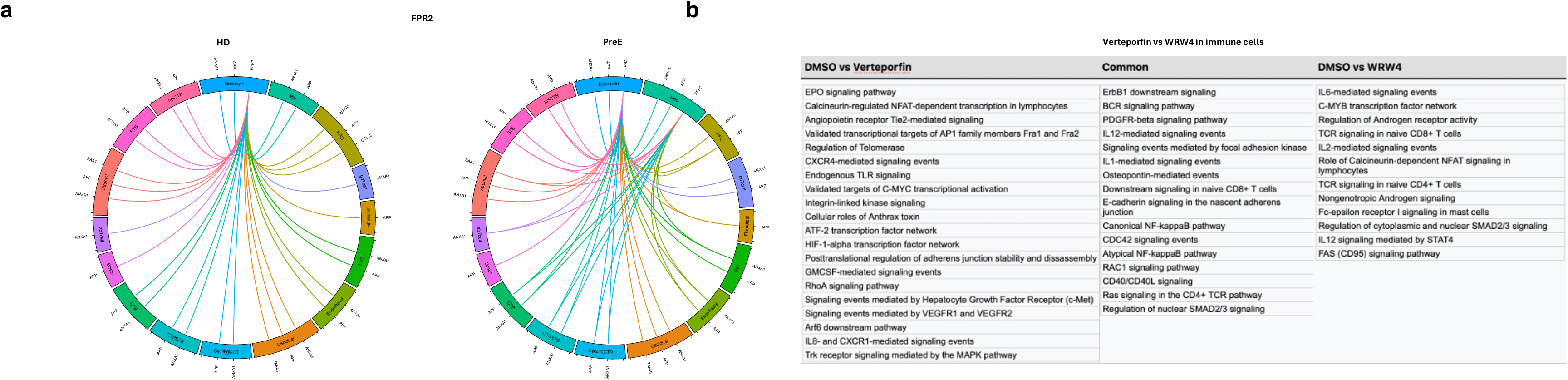
a. FPR2 interaction with its ligands in placental cells from healthy donors and PreE patients. b. Pathways that are unique and common to Verteporfin and WRW4, a FPR2 antagonist.

## STAR METHODS

### Human placentas

The University of Michigan Institutional Review Board has approved collection of human placentas for this study (IRB HUM00184982). All patients gave written consents. Patient demographics data is listed in Table 1. PreE severe features were defined as per the American College of Obstetricians and Gynecologists (ACOG) Task Force report on hypertension in pregnancy (50). Immediately following delivery, whole-thickness placental samples and chorioamniotic membranes were digested with Miltenyi Biotec umbilical cord dissociation kit to obtain single-cell suspension as per the manufacturer’s protocol. Briefly, tissues were washed, cut into small (2-3mm) pieces with sterile scissors or blade, incubated with a digestive enzyme cocktail for 2 hours at 37°C, and dissociated with gentleMACS dissociator run on spleen_01 program. For *ex vivo* treatments, PreE placental explants were incubated with inhibitors or DMSO as a negative control at 4°C overnight prior to digestion with the enzyme cocktail. Samples were then run through 100-um strainers, incubated with red blood cell lysis buffer (Biolegend), washed, and resuspended in IMDM media supplemented with 10% FBS for delivery to the UM Advanced Genomics Core for scRNA-seq. Cell viability was confirmed to be above 80% for all samples. Libraries were constructed on the 10X Chromium system with chemistry v3 and sequenced on the Illumina NovaSeq 6000 sequencer to generate 150 bp paired-end reads.

### scRNA-seq analyses

Data was pre-processed in Seurat (51) by removing genes expressed in fewer than 2 cells and excluding cells that were outliers for number of RNA molecules, or more than 10% mitochondrial genes. DoubletFinder (52) was used to identify and remove probable doublet cells (PC 1:15, PN default 0.25, nExp determined with estimated doublet based on loading density for 10X platform) and DecontX (53) was used to remove probable RNA contamination from individual cells. The individual samples were merged into one object, and pre-processed with the standard Seurat analysis including normalization, variable feature identification, data scaling, and principal component analysis (PCA). The samples were integrated using the *RunHarmony* (10) command in Seurat to mitigate batch effects. Uniform Manifold Approximation and Projection (UMAP) dimensional reduction was done using the *RunUMAP* function in Seurat. Unsupervised clustering was performed by identifying the nearest neighbors using the first twenty dimensions and identifying clusters (resolutions used: human untreated: 0.1, human inhibitor treated: 0.5, mouse Vgll3 OE: 0.3, mouse LPS treated: 0.3). The top cluster-defining genes were found using the *FindAllMarkers* function (absolute log2 fold change threshold of 0.5), and these cluster-defining genes were used to annotate the cell types based on canonical gene signatures and previously published data (54). PE specific genes were identified by using FindMarkers to identify differentially expressed genes between HD and PE on total placenta with a minimum Log2 fold change threshold of +0.1. Putative upstream regulators of this gene signature were identified using Ingenuity Pathway Analysis and plotted on a volcano plot showing the activation z-score and FDR. The proportion of *VGLL3* expressing cells was calculated by taking the number of *VGLL3*+ cells in each individual cluster divided by the total number of cells within the dataset. To perform higher resolution analysis of endothelial and trophoblast clusters, EVT, EVT2, VSM, ImmEC, LymphEC, and CapEC were subsetted into a Seurat object and re-processed including normalization, variable feature identification, data scaling, and PCA identification. Clustering was performed with a resolution of 0.2.

### Gene signature definition and scoring

The composition of the Hippo pathway related genes and fibrosis signature are in Sup. Table 4. PE specific gene signatures were identified with the *FindMarker* command on individual clusters with a minimum Log2 fold change threshold of +0.1 and p-value < 0.05. The *AddModuleScore* function was used to score individual cells for these signatures (ctrl = 10). To score PE gene signatures on mouse data, mouse orthologs of signatures were identified and scored. *Vgll3*-OE specific genes were identified with the *FindMarker* command on individual clusters with a minimum Log2 fold change threshold of +0.1, minimum percentage expressed of 10%, and a p-value < 0.05. Human orthologs of these signatures were scored on the human dataset with the AddModuleScore function.

### Pseudotime trajectory construction

Trophoblast clusters (CTB, transitional trophoblasts, STB, and EVT) were subsetted into a Seurat object and separated by disease state. These objects were imported into Monocle3 by generating a cell data set from the decontaminated counts slot. Normalization and PCA were done with the *preprocess*_*cds* command from Monocle3 with the first 100 dimensions using log normalization, and batch correction was performed using the *align*_*cds* command with continuous effects, which uses the Batchelor tool (55). UMAP dimensional reduction was performed using the *reduce*_*dimension* command. Cells were clustered with the *cluster_cells* command with default parameters. The trajectory graph was learned on the Monocle-derived clusters with *learn*_*graph*. Cells on the UMAP plot are colored by Seurat-derived clusters. Pseudotime was determined using CTB as the starting point. Genes that increase with pseudotime in either the STB or EVT lineage were identified using the *find_gene_modules* function and selecting the modules that increased expression in either lineage. These gene lists were then filtered for those genes with higher expression in PE than HD cells. Ingenuity Pathway Analysis was then used to identify putative upstream regulators of these signatures.

### Ligand-receptor interaction analyses

CellphoneDB (v4.1.0) was used to identify putative ligand-receptor interactions from scRNA-seq datasets (56). To identify interactions between VGLL3+ EVT cells and other cell types in PreE placenta, a new cell-identity was created to denote cells that were positive or negative for VGLL3 expression. Interactions with p < 0.05 were retained, and the number of interactions was plotted with ktplots on a heatmap and on a barchart. Select interactions were plotted on dotplots showing the strength of the interaction. Mouse *Vgll3*-OE and LPS-treated control vs *Vgll3* KO analyses were performed on each genotype separately, and then select interactions were shown on dot plots. Inhibitor-treated human samples were run separately, and significant interactions retained. A summation of all interactions was generated and plotted as a heatmap for each condition. Ligand-receptor interactions that were mutually expressed within the same cell types between the conditions were removed and the resulting non-overlapping interactions were also plotted as a heatmap. Ligands and receptors that were associated with a cytokine, complement, or Hippo pathway were plotted as Circos plots, also removing mutual interactions between conditions.

### Spatial sequencing

FFPE placentas from a PreE patient and HD were re-imbedded into a single block, sectioned unto a slide, and subjected to Xenium in situ transcriptomics preparation and sequencing at UM Advanced Genomics Core. For data analyses, we initially employed *Symphony* (57) to annotate our spatial Xenium dataset by mapping it onto a well-curated, annotated single-cell RNA-seq reference obtained from our current work. This approach enabled robust and accurate cell type identification. Subsequently, we utilized the R package *Giotto* (58) to generate *in situ* plots that visually represent both the cell type annotations and the spatial expression patterns of targeted genes. Finally, differential gene expression analysis was performed using *scran* (59) as implemented within the Giotto framework, facilitating the robust identification of gene expression differences across distinct cellular populations.

### Double-axis scatterplots

DEGs significantly changed (p<0.05) between treatment and control groups for each cell type of interest were inputted into EnrichR to assess pathway enrichment. The compiled pathway data was presented to AI to automate identification of the pathways shared among all cell types, with at least top 3 pathways for each cell type chosen for representation. The double-axis scatterplots were created in Tableau with “Cell Types” as columns and “Pathways” as rows and the intersection of these two variables as the negative log p-values with larger (more significant) values corresponding to heavier dots in the scatterplot.

### Bulk RNA-seq

Five 20-um sections per sample were obtained from FFPE biopsies. RNA isolation was performed using the Qiagen RNeasy FFPE Kit (73504). In brief, sections were transferred to a microcentrifuge tube treated with Qiagen deparaffinization solution and briefly incubated at 56°C. Following a subsequent incubation in a proteinase K lysis buffer at 56°C, tissue sections were shifted to a higher temperature (80°C) to partially reverse formalin crosslinking of the released nucleic acids. This was followed by DNase treatment optimized to eliminate all genomic DNA. The obtained lysates were mixed with Buffer RBC and ethanol and subjected to an RNeasy MinElute spin column. The obtained RNA was eluted with 30 µl of RNase-free water. Libraries were prepared using the QuantSeq 3’ mRNA-Seq Library Prep kit and sequenced on the Illumina NovaSeq 6000 SP Flow Cells. For RNA-Seq analyses, adapter trimming and quality control were conducted on the raw sequence reads. The paired-end reads were mapped using STAR (60) to Genome Reference Consortium Human Build 37 (GRCh37). Only uniquely mapped reads were used for subsequent analysis. Gene expression levels were quantified with GENCODE v24 used as a reference and normalized by HTSeq (61) and DESeq2 (62).

### IHC/ PLA

Formalin-fixed, paraffin-embedded human and mouse tissue sections were heated at 60°C for 30 minutes, deparaffinized, and rehydrated. Slides were placed in either pH9 or pH6 antigen retrieval buffer dependent on the antibody specification. Slides were cooled in ice, treated with 3% H_2_O_2_ for 5 minutes, and blocked with 10% serum for 30 minutes. Primary antibodies included the following: NE (Abcam, ab310335), CD3 (Abcam, ab215212), CD19 (Thermofisher, pa5-27442), VGLL3 (Sigma, HPA054983), FLT1 (Abcam, ab2350), TEAD1 (Santa Cruz, sc-393976), TEAD2 (LS Bio, LSIZC342577), TEAD3 (Creative Biolabs, CBMAB-0234-LY), TEAD4 (Santa Cruz, sc-390578). After an overnight incubation at 4°C, sections were washed and incubated with secondary antibody for 1 hour at room temperature. Following counterstaining with hematoxylin and bluing reagent, slides were imaged under a Zeiss Axioskop 2 microscope. For PLA, five-micron sections were prepared from formalin-fixed, paraffin-embedded human placenta samples and stained using Duolink® Proximity Ligation Assay kit according to the manufacturer’s instructions (Sigma Aldrich).

### Cell Culture, transfections, IP, and protein analyses

HTR8 cells were cultured in Gibco RPMI 1640 (Invitrogen) with 10% fetal bovine serum. HTR8 cells were grown to over 70% confluence and then transfected using Lipofectamine 3000 (ThermoFisher) according to the manufacturer’s protocol. For IP, 500ug-1mg total protein was incubated overnight with anti-FLAG antibody (Origene, TA50011) on a shaker at 4°C. Protein A agarose was used to pull down immune complexes, which were then analyzed by immunoblotting and mass spectrometry. For immunoblotting, proteins were denatured in sample buffer containing β-mercaptoethanol and boiled at 100 °C for 3 min. Samples were separated on gradient 4–20% SDS-PAGE gels (ThermoFisher) and electroblotted to nitrocellulose. Primary antibodies included the following: TEAD1 (Origene, RG215492), GFP (Origene, TA150041), FLAG (Origene, TA50011), VGLL3 (Sigma, HPA054983). Bands were detected on the iBright system using ECL substrate. LC-tandem mass spectrometry was performed at UM Proteomics Resource Facility.

#### Design and validation of Cre-inducible *Vgll3* mice

The *Rosa26* locus on chromosome 6, a “safe harbor” genomic site, was used for the integration of mouse *Vgll3* downstream of a floxed EGFP with STOP sequence to achieve conditional gene expression in mice. Mouse *Vgll3* with *IRES* DNA (1560 bp) was PCR-amplified from *K5-Vgll3-IRES-mCherry*, the construct used for generating our previous K5 promoter-driven *Vgll3* transgenic mouse model (20). The PCR fragment was then cloned into a previously generated *CAG-loxP-EGFP-STOP-loxP-rtTA3-P2A-mCherry-SV40-polyA* cassette, where it replaced *rtTA3-P2A*. This cassette was built by replacing the sequence from CAG up to the *Rosa26* right arm in the backbone of pR26 CAG AsiSI/MluI plasmid (63) (Addgene, 74286). We designed cloning protocols that were carried out by GenScript. The 15.5-kb plasmid was knocked into the mouse *Rosa26* locus of fertilized oocytes using the CRISPR-Cas9 system by the UM Transgenic Animal Model Core. Three transgenic mouse lines were validated by analyzing high molecular weight genomic DNAs extracted from F1 pups with long range PCR (NEB# M0323S LongAmp *Tag* DNA polymerase), to confirm proper 5’ and 3’ integration into the *Rosa26* locus. 2058 bp of 5’ flanking sequences were PCR amplified using primer pair of 5’CTAGGTAGGGGATCGGGACTCTG and 5’AGTAGGAAAGTCCCATAAGGTC (targeting CMV promoter). 5493 bp of 3’ flanking sequences were PCR amplified using primer pair of 5’CACCATCGTGGAACAGTACGAAC (targeting mCherry) and 5’ CAGTGGCTCAACAACACTTGGTC.

### Mouse breeding and phenotyping

*Cyp19-Cre* males were time-mated with *R26-LSL-Vgll3* females to induce *Vgll3* overexpression specifically in the placenta. Detection of a vaginal plug in the morning following coitus was designated as embryonic day (E) 0.5. Blood pressure monitoring was performed at E16.5 using a tail-cuff method on CODA Noninvasive Blood Pressure Monitoring System (Kent Scientific) at UM Physiology Phenotyping Core. Genotyping was carried out by Transnetyx automated service using custom designed probes. Placental histological examinations were conducted by a pathologist on 10 hematoxylin and eosin-stained sections of murine placentas at the Unit for Laboratory Animal Medicine (ULAM) Pathology Core, where complete blood count (CBC) and serum chemistries were also performed. PlGF in maternal circulation was measured by ELISA as per the manufacturer’s protocol (LS Bio, LS-F5507-1). Ejection fraction was measured at E15.5 during ultrasound scanning with Echo at UM Physiology Phenotypic Core. For LPS administration (Sigma-Aldrich, L2630), intraperitoneal injections were performed daily for 10 days starting at E7.5 as described previously (64), and placentas were harvested at E18.5. *Vgll3* KO mice (strain ID EM:11403, C57BL/6NTac-Vgll3/H) were obtained from MRC Harwell Institute. They were generated by CRISPR-induced deletion of 890nt from *Vgll3* gene, including a critical exon 2 ENSMUSE00000876808, to induce a premature stop codon and a null allele.

### Animal body composition analysis

Body fat, lean mass, free water, and total water were quantified using an NMR-based EchoMRI™ 4in1-500 analyzer. Conscious mice were placed individually in a clear plastic holder and inserted into the analyzer without anesthesia or sedation, allowing for non-invasive, rapid measurements within two minutes. The EchoMRI system differentiates tissue composition by exploiting differences in relaxation times of hydrogen proton spins, generating high-contrast signals through specialized radio pulse sequences. Calibration was performed daily using a canola oil reference sample as per manufacturer recommendations.

## EXCEL TABLE TITLES AND LEGENDS

**Supplemental Table 1: Cluster defining genes for human scRNA-Seq.**

**Supplemental Table 2: VGLL3 IP/ mass spectrometry in HEK293 cells.**

**Supplemental Table 3: Cluster defining genes for mouse scRNA-Seq.**

**Supplemental Table 4: Genes used for module scores.**

## REFERENCES

1. Chappell LC, Cluver CA, Kingdom J, Tong S. Pre-eclampsia. Lancet. 2021;398(10297):341–54.

2. Than NG, Romero R, Posta M, Gyorffy D, Szalai G, Rossi SW, et al. Classification of preeclampsia according to molecular clusters with the goal of achieving personalized prevention. J Reprod Immunol. 2024;161:104172.

3. Maynard SE, Min JY, Merchan J, Lim KH, Li J, Mondal S, et al. Excess placental soluble fms-like tyrosine kinase 1 (sFlt1) may contribute to endothelial dysfunction, hypertension, and proteinuria in preeclampsia. J Clin Invest. 2003;111(5):649–58.

4. Zeisler H, Llurba E, Chantraine F, Vatish M, Staff AC, Sennstrom M, et al. Predictive Value of the sFlt-1:PlGF Ratio in Women with Suspected Preeclampsia. N Engl J Med. 2016;374(1):13–22.

5. Turanov AA, Lo A, Hassler MR, Makris A, Ashar-Patel A, Alterman JF, et al. RNAi modulation of placental sFLT1 for the treatment of preeclampsia. Nat Biotechnol. 2018.

6. Simard JF, Arkema EV, Nguyen C, Svenungsson E, Wikstrom AK, Palmsten K, et al. Early-onset Preeclampsia in Lupus Pregnancy. Paediatr Perinat Epidemiol. 2017;31(1):29–36.

7. Kamper-Jorgensen M, Gammill HS, Nelson JL. Preeclampsia and scleroderma: a prospective nationwide analysis. Acta Obstet Gynecol Scand. 2018;97(5):587–90.

8. Cheng X, Jiang Y, Chen X, Huang C, Li S. Early age at menarche is associated with an increased risk of preeclampsia and adverse neonatal outcomes: a 6IZyear retrospective study. Arch Gynecol Obstet. 2024;310(2):807–15.

9. An H, Liu X, Li Z, Zhang L, Zhang Y, Liu J, et al. Association of age at menarche with gestational hypertension and preeclampsia: A large prospective cohort in China. J Clin Hypertens (Greenwich). 2023;25(11):993–1000.

10. Korsunsky I, Millard N, Fan J, Slowikowski K, Zhang F, Wei K, et al. Fast, sensitive and accurate integration of single-cell data with Harmony. Nat Methods. 2019;16(12):1289–96.

11. Deer E, Herrock O, Campbell N, Cornelius D, Fitzgerald S, Amaral LM, et al. The role of immune cells and mediators in preeclampsia. Nat Rev Nephrol. 2023;19(4):257–70.

12. Ullah A, Zhao J, Singla RK, Shen B. Pathophysiological impact of CXC and CX3CL1 chemokines in preeclampsia and gestational diabetes mellitus. Front Cell Dev Biol. 2023;11:1272536.

13. Li J, Liu M, Zong J, Tan P, Wang J, Wang X, et al. Genetic variations in IL1A and IL1RN are associated with the risk of preeclampsia in Chinese Han population. Sci Rep. 2014;4:5250.

14. Than NG, Romero R, Fitzgerald W, Gudicha DW, Gomez-Lopez N, Posta M, et al. Proteomic Profiles of Maternal Plasma Extracellular Vesicles for Prediction of Preeclampsia. Am J Reprod Immunol. 2024;92(4):e13928.

15. Wang X, Yip KC, He A, Tang J, Liu S, Yan R, et al. Plasma Olink Proteomics Identifies CCL20 as a Novel Predictive and Diagnostic Inflammatory Marker for Preeclampsia. J Proteome Res. 2022;21(12):2998–3006.

16. Hosking SL, Moldenhauer LM, Tran HM, Chan HY, Groome HM, Lovell EA, et al. Treg cells promote decidual vascular remodeling and modulate uterine NK cells in pregnant mice. JCI Insight. 2024;10(2).

17. Elsner RA, Smita S, Shlomchik MJ. IL-12 induces a B cell-intrinsic IL-12/IFNgamma feed-forward loop promoting extrafollicular B cell responses. Nat Immunol. 2024;25(7):1283–95.

18. Jang JW, Thuy PX, Lee JW, Moon EY. CXCR4 promotes B cell viability by the cooperation of nuclear factor (erythroid-derived 2)-like 2 and hypoxia-inducible factor-1alpha under hypoxic conditions. Cell Death Dis. 2021;12(4):330.

19. Dey A, Varelas X, Guan KL. Targeting the Hippo pathway in cancer, fibrosis, wound healing and regenerative medicine. Nat Rev Drug Discov. 2020;19(7):480–94.

20. Billi AC, Gharaee-Kermani M, Fullmer J, Tsoi LC, Hill BD, Gruszka D, et al. The female-biased factor VGLL3 drives cutaneous and systemic autoimmunity. JCI Insight. 2019;4(8).

21. Liang Y, Tsoi LC, Xing X, Beamer MA, Swindell WR, Sarkar MK, et al. A gene network regulated by the transcription factor VGLL3 as a promoter of sex-biased autoimmune diseases. Nat Immunol. 2017;18(2):152–60.

22. Hesson AM, Langen ES, Plazyo O, Gudjonsson JE, Ganesh SK. Placental transcriptome analysis of hypertensive pregnancies identifies distinct gene expression profiles of preeclampsia superimposed on chronic hypertension. BMC Med Genomics. 2023;16(1):91.

23. Peng X, He D, Peng R, Feng J, Chen D, Xie H, et al. Associations between IGFBP1 gene polymorphisms and the risk of preeclampsia and fetal growth restriction. Hypertens Res. 2023;46(9):2070–84.

24. Donker RB, Mouillet JF, Nelson DM, Sadovsky Y. The expression of Argonaute2 and related microRNA biogenesis proteins in normal and hypoxic trophoblasts. Mol Hum Reprod. 2007;13(4):273–9.

25. Mouillet JF, Yan X, Ou Q, Jin L, Muglia LJ, Crawford PA, et al. DEAD-box protein-103 (DP103, Ddx20) is essential for early embryonic development and modulates ovarian morphology and function. Endocrinology. 2008;149(5):2168–75.

26. Meinhardt G, Haider S, Kunihs V, Saleh L, Pollheimer J, Fiala C, et al. Pivotal role of the transcriptional co-activator YAP in trophoblast stemness of the developing human placenta. Proc Natl Acad Sci U S A. 2020;117(24):13562–70.

27. Trapnell C, Cacchiarelli D, Grimsby J, Pokharel P, Li S, Morse M, et al. The dynamics and regulators of cell fate decisions are revealed by pseudotemporal ordering of single cells. Nat Biotechnol. 2014;32(4):381–6.

28. Wenzel PL, Leone G. Expression of Cre recombinase in early diploid trophoblast cells of the mouse placenta. Genesis. 2007;45(3):129–34.

29. Chen S, Cao P, Dong N, Peng J, Zhang C, Wang H, et al. PCSK6-mediated corin activation is essential for normal blood pressure. Nat Med. 2015;21(9):1048–53.

30. Fan M, Li X, Gao X, Dong L, Xin G, Chen L, et al. LPS Induces Preeclampsia-Like Phenotype in Rats and HTR8/SVneo Cells Dysfunction Through TLR4/p38 MAPK Pathway. Front Physiol. 2019;10:1030.

31. Wei L, Ma X, Hou Y, Zhao T, Sun R, Qiu C, et al. Verteporfin reverses progestin resistance through YAP/TAZ-PI3K-Akt pathway in endometrial carcinoma. Cell Death Discov. 2023;9(1):30.

32. Verta JP, Moustakas-Verho JE, Donner I, Frapin M, Ruokolainen A, Debes PV, et al. A complex mechanism translating variation of a simple genetic architecture into alternative life histories. Proc Natl Acad Sci U S A. 2024;121(48):e2402386121.

33. Li S, Li A, Zhai L, Sun Y, Yu L, Fang Z, et al. Suppression of FPR2 expression inhibits inflammation in preeclampsia by improving the biological functions of trophoblast via NF-kappaB pathway. J Assist Reprod Genet. 2022;39(1):239–50.

34. Natri H, Garcia AR, Buetow KH, Trumble BC, Wilson MA. The Pregnancy Pickle: Evolved Immune Compensation Due to Pregnancy Underlies Sex Differences in Human Diseases. Trends Genet. 2019;35(7):478–88.

35. Mor G, Aldo P, Alvero AB. The unique immunological and microbial aspects of pregnancy. Nat Rev Immunol. 2017;17(8):469–82.

36. Fu M, Hu Y, Lan T, Guan KL, Luo T, Luo M. The Hippo signalling pathway and its implications in human health and diseases. Signal Transduct Target Ther. 2022;7(1):376.

37. Ma F, Tsou PS, Gharaee-Kermani M, Plazyo O, Xing X, Kirma J, et al. Systems-based identification of the Hippo pathway for promoting fibrotic mesenchymal differentiation in systemic sclerosis. Nat Commun. 2024;15(1):210.

38. Ray S, Saha A, Ghosh A, Roy N, Kumar RP, Meinhardt G, et al. Hippo signaling cofactor, WWTR1, at the crossroads of human trophoblast progenitor self-renewal and differentiation. Proc Natl Acad Sci U S A. 2022;119(36):e2204069119.

39. Hermann A, Wu G, Nedvetsky PI, Brucher VC, Egbring C, Bonse J, et al. The Hippo pathway component Wwc2 is a key regulator of embryonic development and angiogenesis in mice. Cell Death Dis. 2021;12(1):117.

40. Gu B, Ferreira LMR, Herrera S, Brown L, Lieberman J, Sherwood RI, et al. The TEA domain transcription factors TEAD1 and TEAD3 and WNT signaling determine HLA-G expression in human extravillous trophoblasts. Proc Natl Acad Sci U S A. 2025;122(12):e2425339122.

41. Lin Q, Cao J, Yu J, Zhu Y, Shen Y, Wang S, et al. YAP-mediated trophoblast dysfunction: the common pathway underlying pregnancy complications. Cell Commun Signal. 2023;21(1):353.

42. Liu R, Wei C, Ma Q, Wang W. Hippo-YAP1 signaling pathway and severe preeclampsia (sPE) in the Chinese population. Pregnancy Hypertens. 2020;19:1–10.

43. Hu M, Zheng Y, Liao J, Wen L, Cheng J, Huang J, et al. miR21 modulates the Hippo signaling pathway via interference with PP2A Bbeta to inhibit trophoblast invasion and cause preeclampsia. Mol Ther Nucleic Acids. 2022;30:143–61.

44. Kjaerner-Semb E, Ayllon F, Kleppe L, Sorhus E, Skaftnesmo K, Furmanek T, et al. Vgll3 and the Hippo pathway are regulated in Sertoli cells upon entry and during puberty in Atlantic salmon testis. Sci Rep. 2018;8(1):1912.

45. Elks CE, Perry JR, Sulem P, Chasman DI, Franceschini N, He C, et al. Thirty new loci for age at menarche identified by a meta-analysis of genome-wide association studies. Nat Genet. 2010;42(12):1077–85.

46. Oliver JE, Silman AJ. Why are women predisposed to autoimmune rheumatic diseases? Arthritis Res Ther. 2009;11(5):252.

47. Hori N, Okada K, Takakura Y, Takano H, Yamaguchi N, Yamaguchi N. Vestigial-like family member 3 (VGLL3), a cofactor for TEAD transcription factors, promotes cancer cell proliferation by activating the Hippo pathway. J Biol Chem. 2020;295(26):8798–807.

48. Yamaguchi N. Multiple Roles of Vestigial-Like Family Members in Tumor Development. Front Oncol. 2020;10:1266.

49. Soundararajan R, Rao AJ. Trophoblast ‘pseudo-tumorigenesis’: significance and contributory factors. Reprod Biol Endocrinol. 2004;2:15.

50. Hypertension in pregnancy. Report of the American College of Obstetricians and Gynecologists’ Task Force on Hypertension in Pregnancy. Obstet Gynecol. 2013;122(5):1122–31.

51. Stuart T, Butler A, Hoffman P, Hafemeister C, Papalexi E, Mauck WM, 3rd, et al. Comprehensive Integration of Single-Cell Data. Cell. 2019;177(7):1888–902 e21.

52. McGinnis CS, Murrow LM, Gartner ZJ. DoubletFinder: Doublet Detection in Single-Cell RNA Sequencing Data Using Artificial Nearest Neighbors. Cell Syst. 2019;8(4):329–37 e4.

53. Yang S, Corbett SE, Koga Y, Wang Z, Johnson WE, Yajima M, et al. Decontamination of ambient RNA in single-cell RNA-seq with DecontX. Genome Biol. 2020;21(1):57.

54. Vento-Tormo R, Efremova M, Botting RA, Turco MY, Vento-Tormo M, Meyer KB, et al. Single-cell reconstruction of the early maternal-fetal interface in humans. Nature. 2018;563(7731):347–53.

55. Haghverdi L, Lun ATL, Morgan MD, Marioni JC. Batch effects in single-cell RNA-sequencing data are corrected by matching mutual nearest neighbors. Nat Biotechnol. 2018;36(5):421–7.

56. Efremova M, Vento-Tormo M, Teichmann SA, Vento-Tormo R. CellPhoneDB: inferring cell-cell communication from combined expression of multi-subunit ligand-receptor complexes. Nat Protoc. 2020;15(4):1484–506.

57. Kang JB, Nathan A, Weinand K, Zhang F, Millard N, Rumker L, et al. Efficient and precise single-cell reference atlas mapping with Symphony. Nat Commun. 2021;12(1):5890.

58. Dries R, Zhu Q, Dong R, Eng CL, Li H, Liu K, et al. Giotto: a toolbox for integrative analysis and visualization of spatial expression data. Genome Biol. 2021;22(1):78.

59. Lun AT, McCarthy DJ, Marioni JC. A step-by-step workflow for low-level analysis of single-cell RNA-seq data with Bioconductor. F1000Res. 2016;5:2122.

60. Dobin A, Davis CA, Schlesinger F, Drenkow J, Zaleski C, Jha S, et al. STAR: ultrafast universal RNA-seq aligner. Bioinformatics. 2013;29(1):15–21.

61. Anders S, Pyl PT, Huber W. HTSeq--a Python framework to work with high-throughput sequencing data. Bioinformatics. 2015;31(2):166–9.

62. Love MI, Huber W, Anders S. Moderated estimation of fold change and dispersion for RNA-seq data with DESeq2. Genome Biol. 2014;15(12):550.

63. Chu VT, Weber T, Graf R, Sommermann T, Petsch K, Sack U, et al. Efficient generation of Rosa26 knock-in mice using CRISPR/Cas9 in C57BL/6 zygotes. BMC Biotechnol. 2016;16:4.

64. Li G, Ma L, Lin L, Wang YL, Yang H. The intervention effect of aspirin on a lipopolysaccharide-induced preeclampsia-like mouse model by inhibiting the nuclear factor-kappaB pathway. Biol Reprod. 2018;99(2):422–32.

